# The H3K4me1 histone mark recruits DNA repair to functionally constrained genomic regions in plants

**DOI:** 10.1101/2022.05.28.493846

**Authors:** Daniela Quiroz, Diego Lopez-Mateos, Kehan Zhao, Alice Pierce, Lissandro Ortega, Alissza Ali, Pablo Carbonell-Bejerano, Vladimir Yarov-Yarovoy, J. Grey Monroe

**Affiliations:** Department of Plant Sciences, University of California Davis, Davis, CA, USA 95616; Integrative Genetics and Genomics, University of California Davis, Davis, CA, USA 95616; Plant Biology Graduate Group, University of California Davis, Davis, CA, USA 95616; Department of Physiology and Membrane Biology, University of California Davis, Davis, CA, USA 95616; Biophysics Graduate Group, University of California Davis, Davis, California; Institute for Grape and Wine Sciences (ICVV, CSIC-CAR-UR), 26007 Logroño, La Rioja, Spain

**Author notes:** The author(s) responsible for distribution of materials integral to the findings presented in this article in accordance with the policy described in the Instructions for Authors (https://academic.oup.com/plcell/pages/General-Instructions) is (are): Grey Monroe.

## Abstract

Mutation is the ultimate source of genetic variation. Mutation rate variability has been observed within plant genomes, but the underlying mechanisms have been unclear. We previously found that mutations occur less often in functionally constrained regions of the genome in *Arabidopsis thaliana* and that this mutation rate reduction is predicted by H3K4me1, a histone modification found in the gene bodies of actively expressed and evolutionarily conserved genes in plants. We reanalyzed *de novo* germline single base substitutions in fast neutron irradiated mutation accumulation lines in Kitaake rice (*Oryza sativa*) and found the same reduction in mutations associated with H3K4me1, gene bodies, and constrained genes as in *A. thaliana*, suggesting conserved mechanisms for mutation reduction in plants. Here, we characterize a model of targeted DNA repair to explain these observations; PDS5C and MSH6 DNA repair-related proteins target H3K4me1 through their Tudor domains, resulting in nearby DNA experiencing elevated repair. Experimental data and *in-silico* modeling support the high affinity of the Tudor domain for H3K4me1 in both proteins, and that this affinity is conserved between plant species. ChIP-seq data from PDS5C confirms its localization to conserved and low mutation rate genome regions. Somatic and germline mutations observed by deep sequencing of wild-type and *MSH6* knockout lines confirm that MSH6 preferentially repairs gene bodies and H3K4me1-enriched regions. These findings inspire further research to characterize the origins of mechanisms of targeted DNA repair in eukaryotes and their consequences on tuning the evolutionary trajectories of genomes.

## Introduction

Mutations occur when DNA damage or replication error goes unrepaired. Mechanisms that localize DNA repair proteins to certain genome regions, such as by binding certain histone modifications, reduce local mutation rates. Interactions between DNA repair and histone modifications are predicted to evolve if they promote repair in regions prone to deleterious mutations, such as coding regions of essential genes (Lynch, 2010; Martincorena and Luscombe, 2013; Lynch et al., 2016).

Associations between histone modifications and mutation rates have been observed across diverse organisms (Habig et al., 2021; de la Peña et al., 2022; Yang et al., 2021; Yan et al., 2021; Monroe et al., 2022; Makova and Hardison, 2015; Schuster-Böckler and Lehner, 2012). The localization of DNA repair proteins responsible for such mutation biases has been well-established in humans (Supek and Lehner, 2015, 2017, 2019; Foster et al., 2015; Katju et al., 2022). In vertebrates, H3K36me3 is targeted by PWWP domains in proteins contributing to homology-directed and mismatch repair, with H3K36me3 marking the gene bodies and exons of active genes (Li et al., 2013; Huang et al., 2018; Fang et al., 2021; Aymard et al., 2014; Sun et al., 2020). As predicted, reduced mutation rates in active genes and gene bodies have been observed in humans and other animals (Moore et al., 2021; Akdemir et al., 2020; Li et al., 2021; Katju et al., 2022; Supek and Lehner, 2017).

Reduced mutation rates in gene bodies and active genes have also been observed in some algae and land plants, but the mechanism has been unclear (Belfield et al., 2021; Lu et al., 2021; Zhu et al., 2021; Monroe et al., 2022; Belfield et al., 2018; López-Cortegano et al., 2021; Yan et al., 2021; Krasovec et al., 2017). Knockout lines of *MSH2*, the mismatch repair protein that dimerizes with MSH6 to form MutSα (Adé et al., 1999; Kolodner, 1996), indicate that mismatch repair can preferentially target gene bodies in plants. Yet, a specific mechanism of such targeting is unresolved (Belfield et al., 2018).

In plants, H3K4me1 marks gene bodies of active genes. Recent work has demonstrated that H3K4me1 enrichment is mediated by a combination of transcription-coupled (ATXR7) and epigenome-guided (ATX1, ATX2) methyltransferases (Oya et al., 2021). Once established, H3K4me1 reading can occur by proteins containing histone-reader domains such as “Royal family” Tudor domains, which bind methylated lysine residues on H3 histone tails (Kim et al., 2006; Lu and Wang, 2013; Maurer-Stroh et al., 2003).

Recently, the Tudor domain of PDS5C was shown to specifically bind H3K4me1 in *A. thaliana* (Niu et al., 2021). This gene is also a cohesion cofactor that facilitates homology-directed repair (Pradillo et al., 2015; Phipps and Dubrana, 2022; Morales et al., 2020). Recent studies of CRISPR-mediated mutation efficiency show that H3K4me1 is associated with lower mutation efficacy (R = -0.64), supporting more efficient repair (Weiss et al., 2022; Schep et al., 2021; Zhu et al., 2021). These findings are consistent with analyses of mutation accumulation lines in *A. thaliana*, which find H3K4me1 associated with lower mutation rates (Monroe et al., 2022). Still, the extent and consequences of targeted DNA repair remain contentious (Liu and Zhang, 2022).

Here, we reanalyzed *de novo* mutations from whole-genome-sequenced fast neutron mutation accumulation lines in Kitaake Rice (*O. sativa*) (Li et al., 2017) showing similar patterns of mutation reduction and H3K4me1 association as *A. thaliana*. We then characterize a mechanistic model to explain these observations, involving PDS5C and MSH6 proteins targeting H3K4me1 via Tudor domains.

## Results and Discussion

### Mutations in *O. sativa* FN-irradiated lines reflect no selection on SBS mutations

Mutagenesis has been used extensively in the generation and study of plant mutation. With single base substitutions (SBS) in fast-neutron mutation accumulation lines largely reflecting native mutational processes (Wyant et al., 2022; Li et al., 2017), the distribution of these *de novo* mutations could provide insights into mechanisms underlying intragenomic heterogeneity in mutation rate.

We first reanalyzed *de novo* mutations in a population of 1,504 *O. sativa* lines that accumulated mutations upon fast neutron radiation. These data were previously described, and single base-pair substitutions (SBS) were validated with a >99% true positive rate (Li et al., 2017). In total, these data included 43,483 SBS, comprising a combination of indistinguishable fast neutron-related and “spontaneous” mutations (Fig. 1). We restricted our analyses to SBS mutations to preclude selection that may have occurred in the generation of these lines on insertions, deletions, and structural variants (i.e., whole gene deletions). To quantitatively evaluate whether selection on SBS mutations occurred in these lines, we examined non-synonymous and synonymous mutation rates. The ratio of non-synonymous to synonymous mutations in mutation accumulation lines was 2.33 (N/S=5,370/2,155), a 190% increase over this ratio (Pn/Ps = 1.21) observed in polymorphisms of 3,010 sequenced *O. sativa* accessions (Wang et al., 2018) (X^2^ = 670.63, p<2×10^−16^). The ratio of non-synonymous to synonymous *de novo* mutations was not higher in transposable elements (TE) (N/S=2.31) than in non-TE protein-coding genes (N/S=2.34) (X^2^ = 0.035, p = 0.85) nor was it less in coding genes than neutral expectations (N/S=2.33) based on mutation spectra and nucleotide composition of coding regions in the *O. sativa* genome (X^2^ = 0.029, df = 1, p = 0.86) (Fig. S1, methods). To confirm that this non-significance was not simply due to a lack of power, we tested how many non-synonymous mutations would have to be “missing” because of selection to explain the 29% reduction in mutation rate of coding genes relative to non-genic (intergenic, TE) regions we observed in downstream analyses. Given that coding regions constitute only 37% of gene bodies (the rest being intronic and untranslated exonic regions), we find that 34% of non-synonymous mutations would have to have been missing due to selection, which would be readily detected by a significant deviation from the null expectation (X^2^ = 224.4, df = 1, p < 2e-16). In conclusion, we detect no evidence of selection on SBS mutations, providing an opportunity to study *de novo* mutation rate heterogeneity before selection.

**Figure 1.**
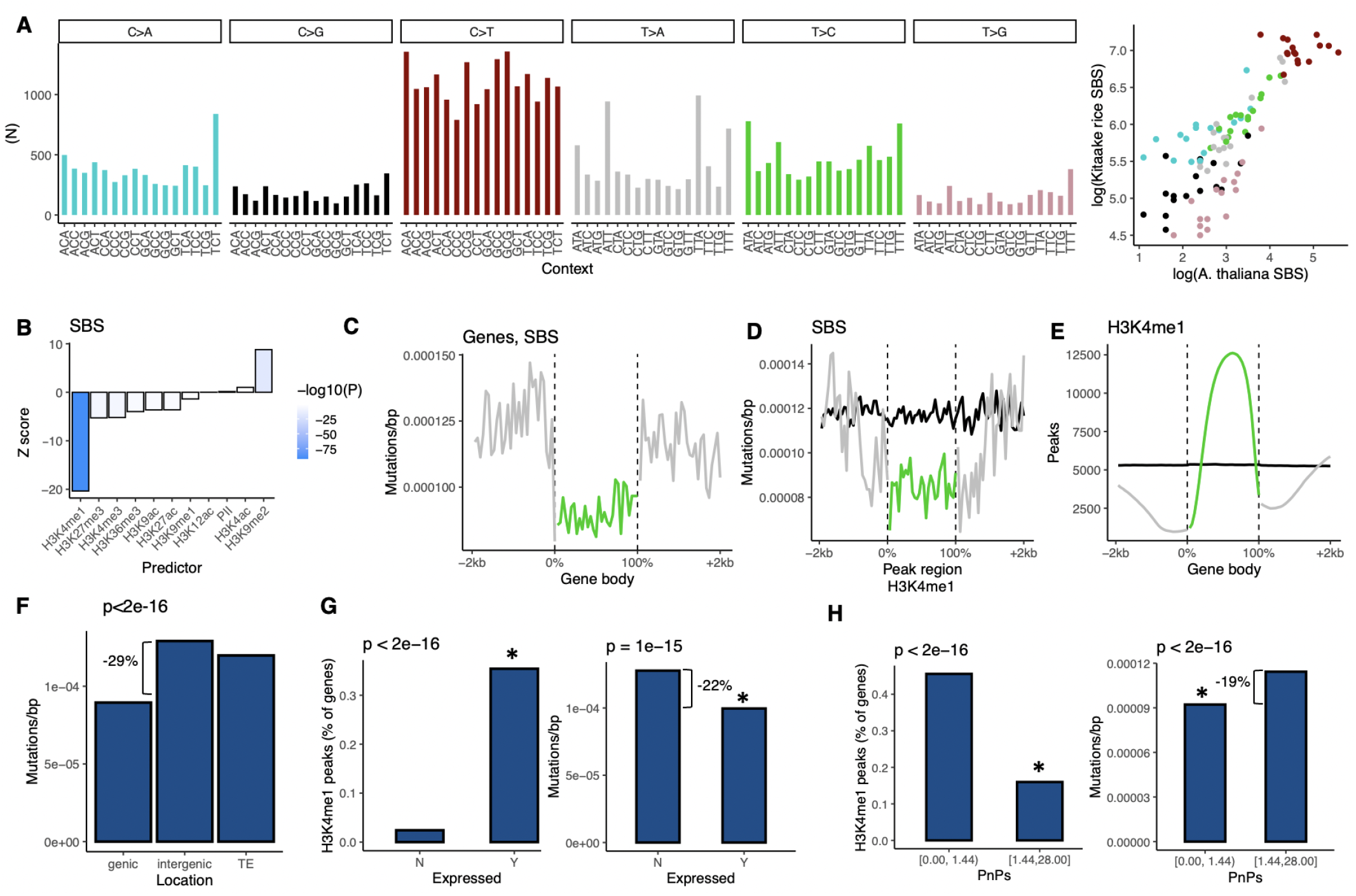
Spectra and distribution of germline SBS mutations in fast neutron rice. **A**, Frequency of mutations in trinucleotide contexts and correlation with *de novo* germline mutations in *A. thaliana* (r>0.8, p<2×10 ^−16^) (Weng et al. 2019, Monroe et al. 2022). **B**, Results from logistic regression modeling mutation probability in 100bp windows as a function of overlap with epigenomic marks around genes in rice. **C**, Mutation rates in relation to gene bodies in rice. D, Mutation rates in relation to H3K4me1 peak regions in rice. **E**, H3K4me1 peaks regions distribution in gene bodies in rice. **D-E**, Black line shows null expectation (mutations around randomized peaks). **F**, Mutation rates in genic, intergenic, and TE regions. **G**, H3K4me1 enrichment and gene body mutation rates in genes annotated as expressed (Y) and non-expressed (N) in rice. **H**, H3K4me1 enrichment and gene body mutation rates in genes subject variable degrees of purifying selection (low vs high Pn/Ps) in 3,010 natural accessions of rice.

We then compared SBS spectra in trinucleotide contexts from these lines with *de novo* germline mutations in *A. thaliana* (Weng et al., 2019; Monroe et al., 2022) and found that they are significantly correlated (Fig. 1A, r=0.8, p<2×10^−16^). We cannot know how much of the residual difference in SBS spectra is due to the effects of fast neutron mutagenesis versus natural differences between *O. sativa* and *A. thaliana*, as variations in the spectra of SBS have been reported between related species and even different genotypes of the same species (Jiang et al., 2021; Sasani et al., 2022; Cagan et al., 2022). Future work will benefit from the evaluation of *de novo* mutations arising in different species under diverse conditions to understand the environmental and genetic controls of SBS mutational spectra in plants.

### H3K4me1 marks low mutation rate regions in *O. sativa*, including gene bodies, and transcriptionally active and evolutionarily conserved genes

To test whether the genome-wide distribution of mutations in *O. sativa* is associated with histone modifications, we used data from the riceENCODE epigenomic database, which includes H3K4me1, H3K9me1, H3K4me3, H3K36me3, H3K9me2, H3K27me3, H3K27ac, H3K4ac, H3K12ac, H3K9ac, and RNA polymerase II (PII) measured by chromatin immunoprecipitation sequencing (ChIP-seq) (Xie et al., 2021). We then tested whether mutation probabilities in 100bp windows in genic regions (the probability of observing a mutation) were predicted by epigenomic features and found a significant reduction in mutation probabilities in windows that overlapped with H3K4me1 peaks (Fig. 1B). These data are consistent with observations in other plant species where lower mutation rates have been observed in regions marked by H3K4me1 (Monroe et al., 2022; Weiss et al., 2022). To further confirm that these patterns were not due to selection, we also restricted our analyses to only those genes in which loss-of-function mutations were found in the population (Li et al., 2017) and observed similar results (Fig. S2). We considered that mutation rate heterogeneity was possibly caused by GC>AT mutations in transposable elements with elevated cytosine methylation in non-genic sequences rather than histone-mediated mutation reduction. Therefore, we restricted our analyses to exclude all such GC>AT mutations and observed similar results. H3K4me1-associated hypomutation was also the same when analyses were restricted to only homozygous mutations (Fig. S2). We further repeated analyses with other regression approaches: single predictor binomial regression, multiple linear regression, Poisson regression, and model selection based on AIC. In all cases, H3K4me1 was found to have the most significant association with reduced mutation rates (Fig. S4)

We calculated mutation rates in genes and their neighboring sequences and observed a significant reduction in mutation rates in gene bodies (Fig. 1C), consistent with lower mutation rates in gene bodies of other plants. Mutation rates were lower both in and around H3K4me1 peaks, which could indicate the action of local recruitment and targeting of DNA repair to H3K4me1 marked sequences including gene bodies (Fig. 1D). That mutation rates were also lower in sequences immediately neighboring H3K4me1 peaks could indicate a spatially distributed effect on mutation around H3K4me1, or the effect of conservative peak calling (Fig. 1D). Only 8.9% of H3K4me1 peaks were found outside of non-TE protein-coding genes. Nevertheless, we could use these instances of non-genic H3K4me1 to test whether the reduction in mutation rates in H3K4me1 peaks was due simply to selection against coding region mutations affecting results. When considering all H3K4me1 peaks, we observed a 20.1% reduction in mutation rates compared to regions within 2kb outside of peaks (X^2^ = 124.38, p < 2.2e-16 -). For non-genic H3K4me1 peaks, we observed the same reduction: -20.2% (X^2^ = 9.88, p = 0.00167). Together, these results suggested a role of H3K4me1 in localized hypomutation, which could explain reduced gene body mutation rates observed since gene bodies are enriched for H3K4me1 (Fig. 1E,F).

We then compared mutation rates between different classes of genes, which proved consistent with the expected effects of increased DNA repair in functionally constrained genes caused by H3K4me1-localized repair. Mutation rates were significantly lower in genes that overlapped with H3K4me1 peaks, including when considering only synonymous mutations (Fig S3). H3K4me1 peaks were enriched in genes annotated as expressed compared with those not expressed (Kawahara et al., 2013) (X^2^ = 2550961, p < 2×10^−16^). And, as predicted by their enrichment for H3K4me1, mutation rates were 22% lower in expressed genes (X^2^ = 63.7, p = 1×10^−15^)(Fig. 1G). This result was confirmed in an analysis restricted to synonymous mutations in coding regions only, rejecting the hypothesis that these results are due to selection (Fig. S3). Indeed, the ratio of non-synonymous to synonymous *de novo* mutations in the data was not different between expressed and non-expressed genes (X^2^ = 0.0007, p = 0.98) (Fig. S3). Comparing genes that exhibit different degrees of selection in 3,010 natural accessions of *O. sativa* (Wang et al., 2018), those under elevated purifying selection with low Pn/Ps (non-synonymous/synonymous polymorphisms), were enriched for H3K4me1 peaks (X^2^ = 8045711, p < 2×10^−16^) and experienced 19% lower mutation rates (X^2^ = 188.5, p < 2×10^−16^) (Fig. 1H). These genes did not have a lower ratio of non-synonymous to synonymous in *de novo* mutations (X^2^ = 0.22, p=0.63) (Fig S3) and the reduction of mutation rates in conserved genes was confirmed by analysis of synonymous mutations only (Fig S3). We, therefore, find that mutation rates are significantly lower in functionally constrained genes in *O. sativa*, which cannot be explained by selection on non-synonymous mutations, consistent with what has been previously shown in *A. thaliana* (Monroe et al., 2022).

### PDS5C targets H3K4me1 and is associated with lower mutation rates

These findings in *O. sativa* are consistent with reports of reduced mutation rates in gene bodies of expressed and constrained genes in *A. thaliana* and other species (Krasovec et al., 2017; Moore et al., 2021; Monroe et al., 2022). While in humans, this is known to be mediated by H3K36me3 targeting by DNA repair genes, our results suggest that H3K4me1 may be a target of DNA repair in plants. To examine this further, we considered genes with known H3K4me1 targeting. PDS5C, a gene belonging to a family of cohesion cofactors that facilitate homology-directed repair (HDR), contains a Tudor domain that was recently discovered to specifically bind H3K4me1 (Niu et al., 2021). Analyses of ChIP-seq data of PDS5C-Flag from *A. thaliana* show PDS5C is targeted to gene bodies (which are enriched for H3K4me1 in both *O. sativa* and *A. thaliana*) (Fig. 2A). We also find that PDS5C is increased in regions of lower germline mutation rates in *A. thaliana*, consistent with recruitment and its function in facilitating DNA repair (Fig. 2B).

**Figure 2.**
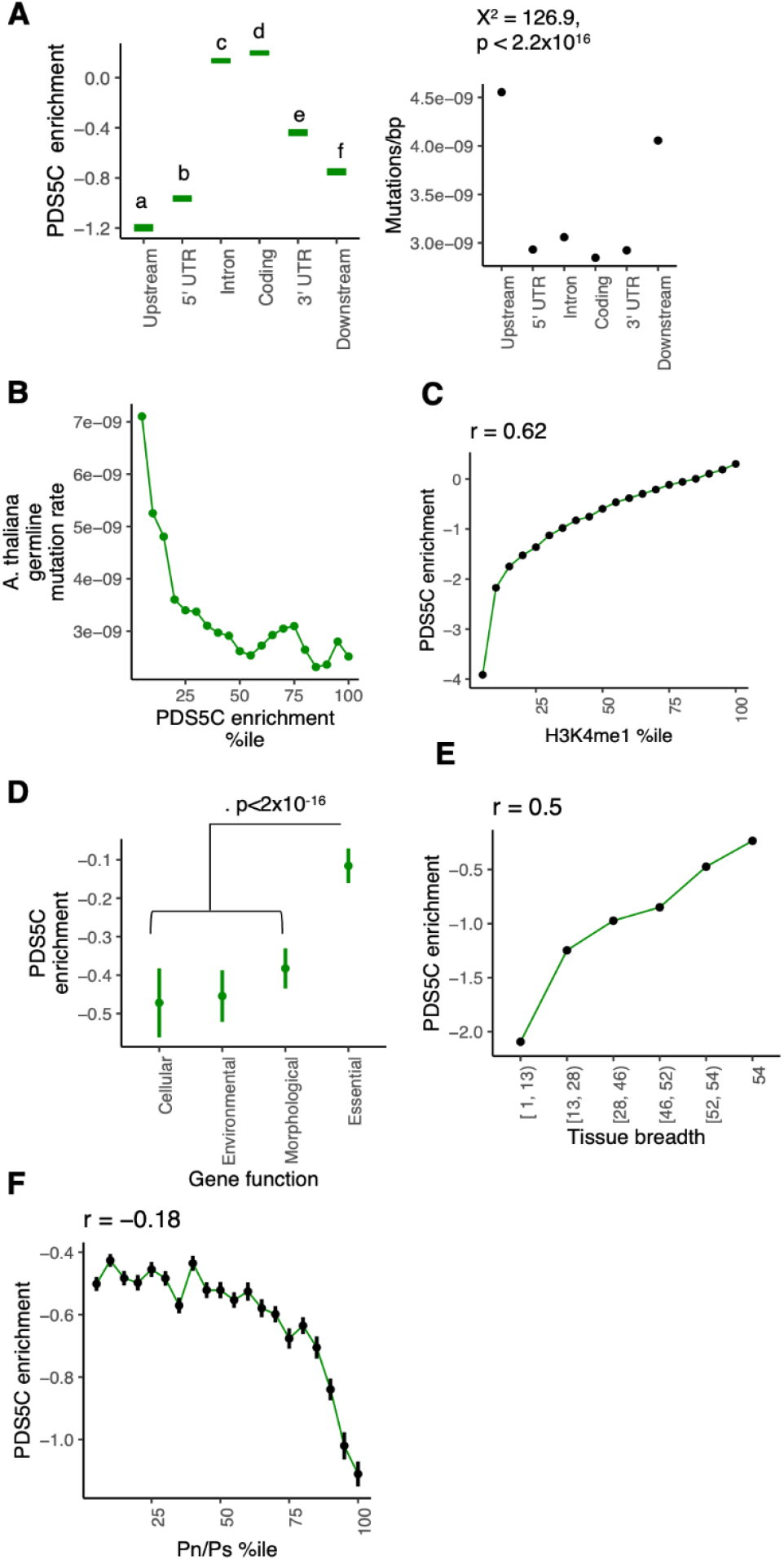
PDS5C targets H3K4me1 and is associated with lower mutation rates. **A**, ChlP-seq analysis of PDS5C in *A thaliana*, cohesion cofactor involved in HDR (Niu et al. 2021). Letters represent TukeyHSD contrast at p=0. Upstream/downstream = 1000bp. **B**, Features marked by PDS5C experience reduced germline mutation rates in *A. thaliana*. **C**, Gene bodies of genes with elevated H3K4me1 have greater levels of PDS5C, as do **D**, essential genes, **E**, genes expressed in many tissues, and **F**, genes under stronger purifying selection. Pn/Ps = Non-synonymous /synonymous polymorphisms in *A. thaliana* populations. Mean +- SE for percentiles shown. Pearson correlation (r) reflects entire dataset. P<2×10^−16^ for all.

Evolutionary models predict that histone-mediated repair mechanisms should evolve if they facilitate lower mutation rates in sequences under purifying selection (Lynch et al., 2016; Martincorena and Luscombe, 2013). As predicted by this theory, we find that PDS5C targeting (ChIP-seq) is enriched in coding sequences, essential genes (determined by experiments of knockout lines (Lloyd and Meinke, 2012; Lloyd et al., 2015)), genes constitutively expressed (detected in 100% of tissues sampled) (Mergner et al., 2020), and genes under stronger purifying selection in natural populations of *A. thaliana* (lower Pn/Ps) (1001 Genomes Consortium, 2016) (Fig. 2C-F).

Visualizing the *A. thaliana* PDS5C protein full-length model generated by Alphafold (Jumper et al., 2021) reveals that the PDS5C active domain is separated from the Tudor domain by unstructured, and potentially flexible segments (Fig. 3A), suggesting that the Tudor domain operates as an anchor, localizing PDS5C to H3K4me1 and gene bodies of active genes. PDS5C is a cohesion cofactor linked to multiple DNA repair pathways. In its role in cohesion between sister chromatids, it has been reported to promote HDR (Pradillo et al., 2015). This is consistent with its known interaction with repair proteins along with the direct contribution of cohesion and PDS orthologs to the HDR pathway (Morales et al., 2020; Phipps and Dubrana, 2022; Hill et al., 2016; Ren et al., 2005; Bolaños-Villegas et al., 2013; Schubert et al., 2009). The observation that mutation rates are reduced at H3K4me1 peak regions (Fig. 1) supports the hypothesis that Tudor domain-mediated targeting in PDS5C, its orthologs (PDS5A, B, D, and E), or other repair-related proteins contribute to targeted hypomutation in the functionally important regions of the genome. Still, additional experiments are needed to quantify the precise local effect of PDS5C on mutation rate. We compared the PDS5C Tudor domain sequence between *A. thaliana* and *O. sativa* and found that the critical amino acids constituting the aromatic cage, where H3K4me1 binding specificity is determined, are conserved (Fig. 3C), suggesting a potential role of PDS5C in the mutation biases observed here in *O. sativa* (Fig. 1).

**Figure 3.**
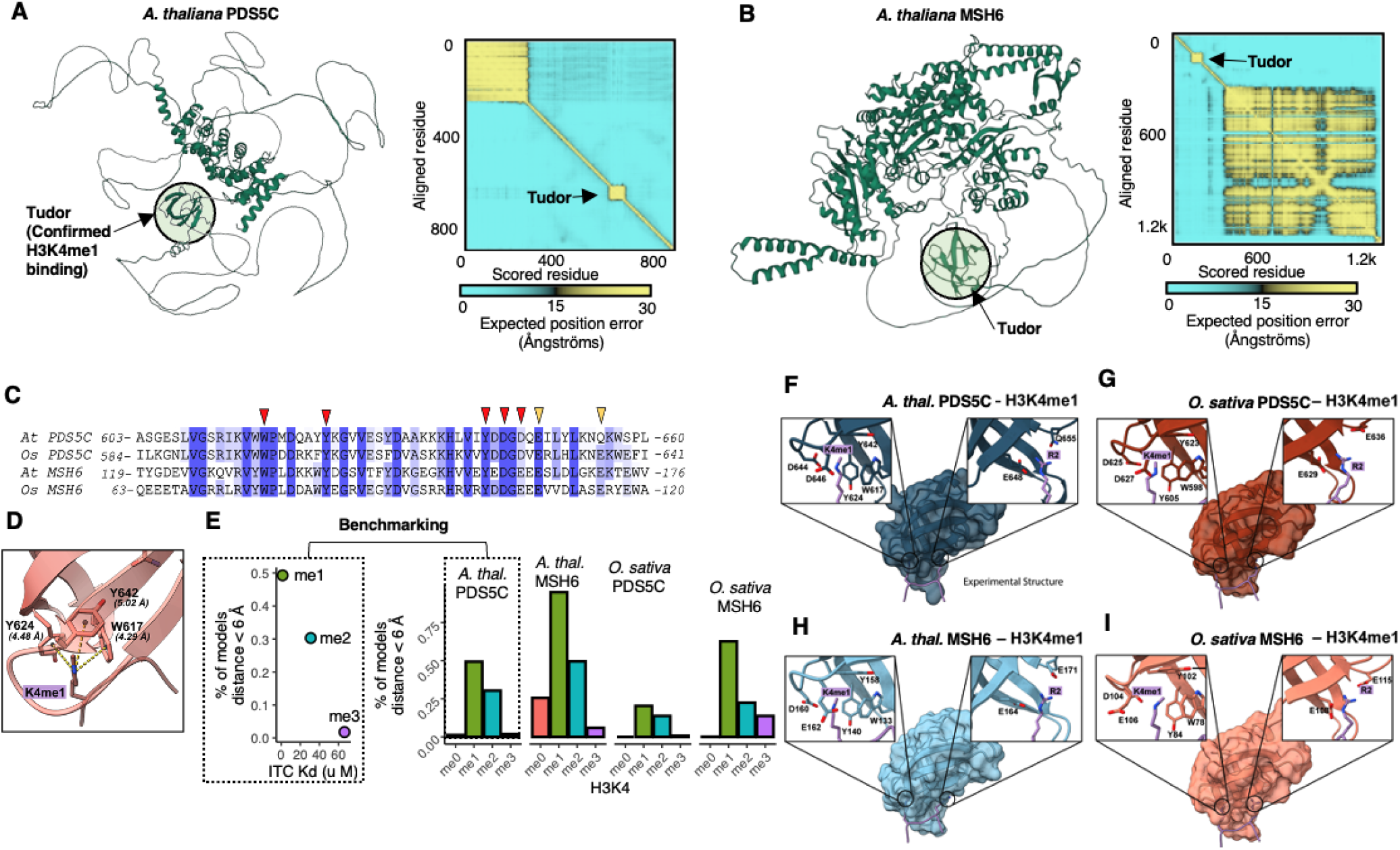
*Arabidopsis thaliana* and *Oryza sativa* PDS5C and MSH6 structure and Tudor domains affinity to H3K4 methylation. **A-B**, Alphafold2 structures of PDS5C and MSH6 protein from *A. thaliana* indicate Tudor domain is tethered to active domains **C**, ClustalW alignment of Tudor domains. Red arrows mark residues shown to be important for H3K4me1 specific binding, and yellow arrows mark residues important to interact with H3R2 (Niu et al. 2021) **D**, Aromatic cage and H3K4me1 in experimental structure of *A. thal*. PDS5CTudor domain (PDB: 7DE9) (Niu et al. 2021) highlighting in yellow the distances K4 nitrogen to the center of the three aromatic rings. **E**, Left panel, correlation between the % of models with K4 within the aromatic cage (distance to aromatic rings < 6Å, see D and Fig S7) and dissociation constant values of H3 tail peptides in different methylation states obtained from ITC experiments for *A. thal*. PDS5C Tudor domain (Niu et al. 2021); right panels, % of models with K4 in each methylation state within the aromatic cages of the Tudor domains of PDS5C and MSH6 from both *A. thaliana* and *O. sativa*. **F-l**, FlexPepDock models of the H3K4me1 peptide in the aromatic cage of Tudor domains of *A. thal*. PDS5C, *A. thal*. MSH6, *O. sativa* PDS5C and *O. sativa* MSH6. Residues interacting with H3K4me1 are amplified in the left boxes. Resides interacting with H3R2 are highlighted in the right boxes.

### Multiple DNA repair mechanisms with Tudor domains could be influencing mutation biases

The discovery of the PDS5C Tudor domain as an H3K4me1 targeting domain (Niu et al., 2021) provides an opportunity to identify other proteins with potential for H3K4me1-mediated gene body recruitment. We used *blastp* to search the *A. thaliana* proteome for other proteins containing Tudor domains similar to that of PDS5C. An analysis of gene ontologies indicated that this gene set is highly enriched for genes with DNA repair functions (9/29 genes, p=1×10^−11^)(Fig. S5A, Table S1). Five of these were PDS5C homologs. We also found that MSH6, a DNA mismatch repair protein, contains a Tudor domain similar to that of PDS5C, which was an obvious candidate for further consideration. That the MSH6 homologous protein in vertebrates instead contains a PWWP that targets gene bodies via H3K36me3 binding suggests a remarkable example of convergent evolution between plants and vertebrates (Li et al., 2013).

Structural modeling prediction of MSH6 structure using AlphaFold (Jumper et al., 2021) indicates that its Tudor domain may, like that in PDS5C, function as an anchor, tethering it to H3K4me1 leading to local increases in DNA repair (Fig. 3A-B, Fig. S5). Sequence comparison suggests that *O. sativa* MSH6, *A. thaliana* MSH6, and *O. sativa* PDS5C Tudor domains may present a similar binding preference for H3K4me1 as *A. thaliana* PDS5C Tudor domain since key residues forming monomethylated lysine binding site are conserved between all homologous domains (Fig. 3C). To test this hypothesis, we modeled *A. thaliana* MSH6 and *O. sativa* MSH6 and PDS5C Tudor domains with AlphaFold and compared them with the experimental structure of *A. thaliana* PDS5C Tudor domain (PDB:7DE9) (Niu et al., 2021). Superimposition of the three modeled Tudor domains onto the PDS5C Tudor domain showed remarkably similar structures with backbone root mean square deviation (RMSD) values below 1 Å (Fig. S6).

Subsequently, we modified the K4me1 from the H3 tail peptide bound to the PDS5C Tudor domain in ChimeraX (Goddard et al., 2018) to obtain H3K4, H3K4me2, and H3K4me3. Using Rosetta FlexPepDock (Raveh et al., 2010), we modeled the docking of the different methylation states of H3K4 to the Tudor domains of PDS5C and MSH6 from *A. thaliana* and *O. sativa*.

We analyzed the geometry of the binding site in the top 10% of models based on Rosetta total score, by measuring the distances of H3K4 nitrogen to the center of the three aromatic rings forming the aromatic cage in the Tudor domains (Fig 3D). The experimental structure of the PDS5C Tudor domain bound to H3K4me1 (PDB:7DE9) (Niu et al., 2021) provided a reference to analyze the geometry of the aromatic cage. In this structure, the average distance between H3K4 nitrogen and the aromatic rings is 4.6 Å (Fig. 3D). We calculated this average distance in the *in silico* models generated by FlexPepDock and analyzed its distributions per case (Fig. S7).

We found that by setting a distance threshold of 6Å, we were able to accurately calculate the proportion of generated models with the H3K4 inside the aromatic cage. For the *A. thaliana* PDS5C Tudor, the proportion of models with H3K4 inside the aromatic cage was a near-perfect correlation (r=0.999) with the dissociation constant values obtained experimentally by ITC (Figure 3E). Therefore, we reasoned that we could apply this approach to predict the binding preference of the Tudor domains of *A. thaliana* MSH6 and *O. sativa* MSH6 and PDS5C to the different epigenetic marks. For all three cases, we observed that H3K4me1 was the mark sampled more often inside the aromatic cage (Fig. 3E), followed by H3K4me2, and H3K4me3, aligning with the experimental results for the Tudor domain of *A. thaliana* PDS5C. Interestingly, lack of methylation resulted in almost no sampling of geometries within the aromatic cage, which aligns with ITC experimental results that show no detectable binding of H3 tail peptide by the PDS5C Tudor domain when K4 was not methylated (Niu et al., 2021). These results suggest that MSH6 and PDS5C Tudor domains bind to H3K4 with more affinity when it is in the monomethylated state (H3K4me1). While binding to H3K4me2 and me3 is possible, our results suggest a larger energetic barrier to access the aromatic cage for these two states and lower affinity. However, binding experiments must be conducted to obtain dissociation constant values, thus determining the exact selectivity for H3K4me1 over H3K4me2 and H3K4me3 of these Tudor domains.

Based on previous experimental data and our computational analysis, we conclude that MSH6 and PDS5C Tudor domains bind to H3K4 preferentially when it is in the monomethylated state (H3K4me1). Interestingly, *in-silico* predictions suggest different levels of affinity between the Tudor domains, which could lead to differences in their contribution to the H3K4me1-associated mutation rate reduction. The similar binding preference of PDS5C and MSH6 Tudor domains suggests that multiple repair pathways could have evolved H3K4me1-targeting potential in plants, motivating additional experiments and investigations into the evolutionary origins of these mechanisms. Because MSH6 operates as a dimer with MSH2 to form the MutSα complex, which recognizes and repairs small mismatches, its Tudor domain could explain the previous observation that MSH2 preferentially targets gene bodies to reduce mutation rates therein (Belfield et al., 2018). These findings are consistent with extensive work showing that mutation rates can be lower in gene bodies of active and conserved genes, but suggest that these mechanisms are independent of, while functionally analogous to, similar mechanisms known in vertebrates (H3K36me3 targeted repair described in the introduction). Further experiments are needed - the reduced mutation rates in gene bodies in active genes in plants may be explained by multiple mechanisms collectively targeting H3K4me1 or additional histone states via Tudor domains as well as other histone modifications and readers (Davarinejad et al., 2022; Liu et al., 2022).

### *de novo* mutations in *MSH6* knockout lines indicate targeted repair of gene bodies and H3K4me1 marked regions

To experimentally test the effect of MSH6 in targeted DNA repair and their potential consequence in the reduction in mutation rates associated with H3K4me1 and therefore gene bodies, we performed deep sequencing (250x depth coverage) of 3-week old rosettes of *A. thaliana* of 7 *MSH6* knockout line SALK_037557 (*msh6*/*msh6*), 1 *MSH6* knockout line SALK_089638 (*msh6*/*msh6*), 2 heterozygous SALK_089638 lines (*WT*/*msh6*), and 9 wildtypes (*WT*/*WT*) lines (Fig. 4A). Variant caller Strelka2 was used in TUMOR-NORMAL mode to call SBS mutations for each sample following a strict filtering process to call true mutations. In summary, variants were kept if they were called in only 1 sample, PASS for all 16 “NORMAL” comparisons reported by Strelka2 (Kim et al., 2018), the distance to nearest mutation >1000 bp, total read depth >150, and alt read support > 5. Somatic *vs de novo* germline mutations were distinguished based on deviation from expected alternative variant frequencies (X^2^ p<0.001, alt reads < 50%) (Fig. S8B). As expected, the total number of somatic mutations was higher in the *MSH6* knockout lines (mean±SE = 40.25±3.14 per sample) than in wild-type and SALK_089638c heterozygous lines (14.1±2.31 per sample). Mutations from heterozygotes were combined with wild-type samples for further comparisons because heterozygous lines exhibited the wild-type phenotypes (Fig. S8A).

**Figure 4.**
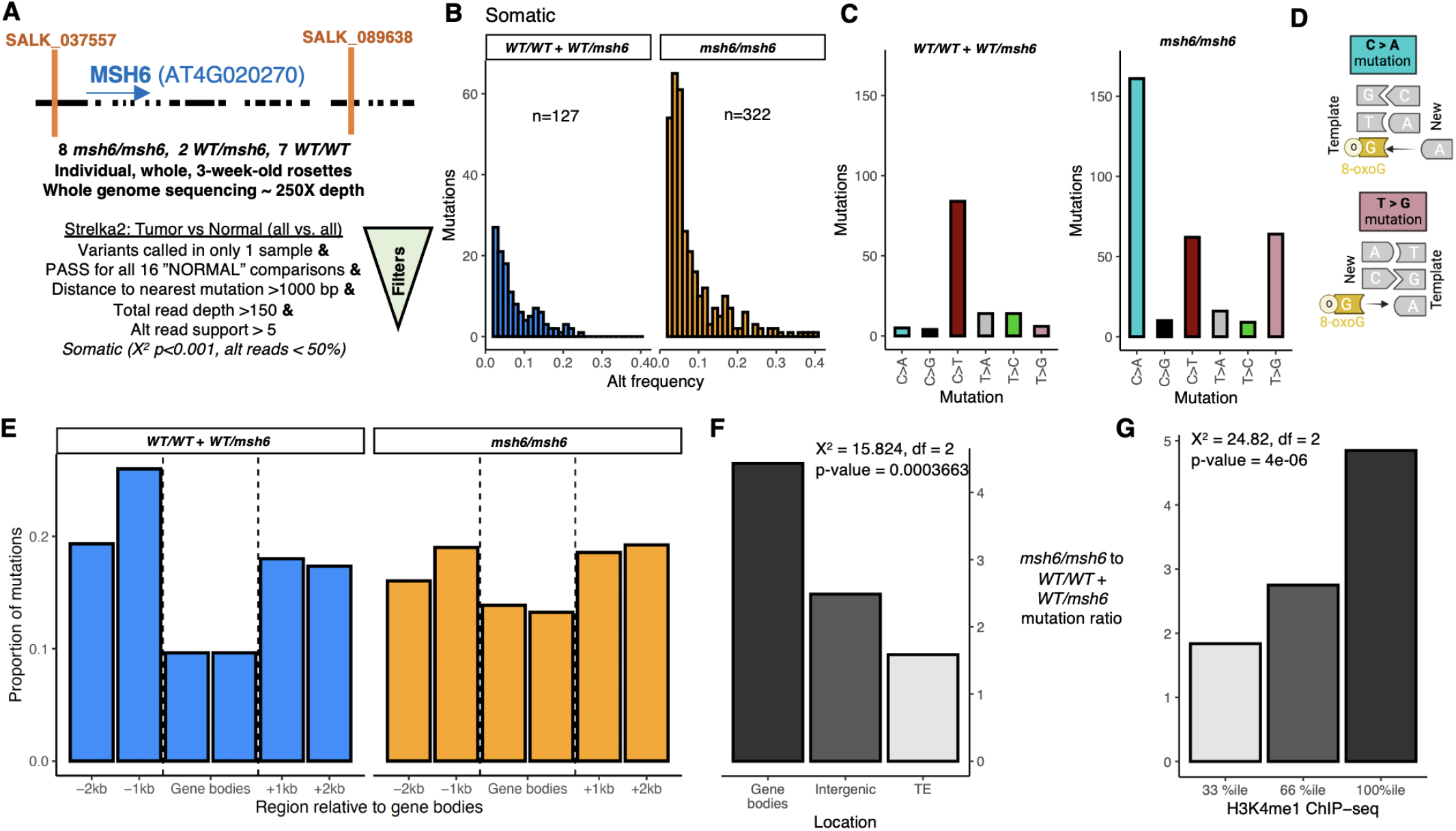
Somatic mutations in MSH6 knockout lines. **A**, Diagram representing *MSH6* mutant SALK insertions used to phenotype mutation rates in *MSH* deficient lines. General pipeline for Somatic mutation experimental design and strict variant calling using Strelka2. **B**, Alternative allele frequency of mutations found in *WT/WT* + *WT/msh6* and *msh6/msh6* lines. The total number of mutations (n) is 127 and 322, respectively. **C**, mutation signature of *Wt/WT* + *WT/msh6* and *msh6/msh6* with *msh6/msh6* showing **D**, expected effects of unrepaired mismatches between adenine and oxidized guanine. **E**, Proportion of mutations distribution in gene bodies in *WT/WT* + *WT/msh6* and *msh6/msh6* lines. **F**, *msh6/msh6* to *WT/WT* + *WT/msh6* mutation ratio in different genomic features and **G**, in genome regions grouped by H3K4me1 ChlP-seq.

As expected due to mismatches arising from the tendency for oxidized guanine (8-oxoG) to mispair with adenine (Fig. 4D) (Kino et al., 2017), C>A and T>G substitutions were proportionally higher in *MSH6* knockout lines (Fig. 4C), which is consistent with mutational signatures observed in mismatch repair (MMR) deficient organisms (Sanders et al., 2021; Lujan et al., 2014). This increase in C>A and T>G SBS was also found for the *de novo* germline mutations (Fig. S6C). This result is consistent with the high affinity of MutSα for 8-oxoG:A mispairings (Mazurek et al., 2002). In light of evidence that MSH6 physically binds MutY (Gu et al., 2002; Hahm et al., 2022) and interacts synergistically with OGG1 (Ni et al., 1999; Pavlov et al., 2003), both proteins responsible for 8-oxoG:A base-excision repair (BER), we cannot exclude the possibility that MSH6 promotes the local activity of multiple (MMR and BER) repair pathways. MSH6 also influences somatic recombination in *A. thaliana* (Li et al., 2006; Gonzalez and Spampinato, 2020). Future studies should also consider the potential effects of somatic recombination on local mutaiton rate.

To test whether MSH6 contributes to the reduction in mutation rates in gene bodies and H3K4me1 marked regions, we explored changes in the proportion of mutations in *MSH6* knockout and wild-type lines. The distribution of mutations in wild-type plants around genes was consistent with what has been previously reported (Monroe et al., 2022), while in *MSH6* knockout mutant lines gene body mutation rates were increased, consistent with MSH6 targeting to gene bodies in wild-type plants (Fig. 4E). In gene bodies, mutation rates were more than 4.5X higher in *MSH6* knockout lines than the wildtype and heterozygous samples, while the increase of mutations in intergenic regions and TEs was significantly less, 2.4X and 1.7X, respectively (Fig. 4F). We also estimated the mutation ratio in relation to H3K4me1 ChIP-seq enrichment, and found that *MSH6* knockout lines contain ∼5X more mutations in genome regions most highly enriched for H3K4me1, with regions containing less H3K4me1 experiencing significantly lower increase mutation rates in the knockout lines (Fig. 4G).

Experimental results are consistent with the hypothesis that H3K4me1 marked genome regions experience targeted DNA repair in plants, with the Tudor domain of repair proteins providing a mechanistic basis for this recruitment. These results inspire future functional studies of this and other potential candidate genes influencing targeted DNA repair and emergent mutation biases.

## Conclusions

We found evidence of mutation bias associated with H3K4me1-mediated DNA repair in *O. sativa* and examined potential mechanisms conserved in plants (Fig. 5). Our observations here are derived from reanalyses of data generated by independent research groups (Li et al., 2017; Niu et al., 2021; Xie et al., 2021) and prove consistent with previous reports of mutation biases in *A. thaliana* (Monroe et al., 2022). The mechanisms revealed are aligned with evolutionary models of evolved mutation bias, indicating targeting of mismatch repair and homology-directed repair pathways to regions of the genome functionally sensitive to mutation: coding regions and genes under stronger evolutionary constraints. Furthermore, we find genetic evidence that MSH6 regulates the reduced mutation rate in gene bodies and H3K4me1-marked regions in *A. thaliana*. Functional characterization of PDS5 proteins is still required to estimate their influence over lower mutation rates in H3K4me1-associated regions. These findings provide a plant-specific and higher-resolution mechanistic model of hypomutation in gene bodies and essential genes, motivating experimental investigations to further elucidate the extent, evolutionary origins, and consequences of chromatin-targeted DNA repair.

**Figure 5.**
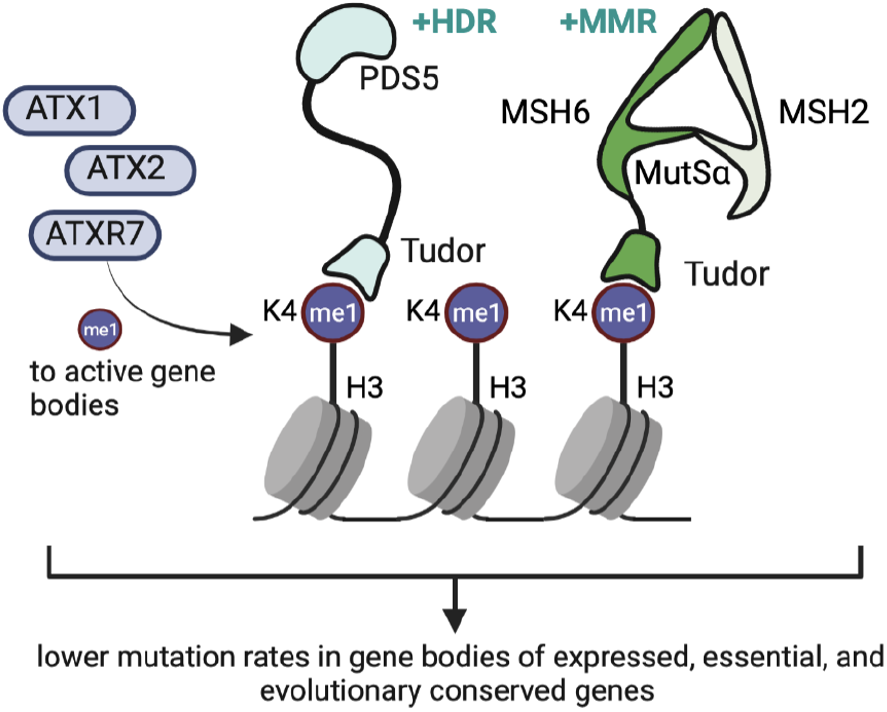
Emerging mechanistic model of H3K4me1-targeted DNA repair in plants. ATX1, ATX2, ATXR7 are responsible for the enrichment of H3K4me1 in the gene bodies of active genes (Oya et al. 2021). Tudor domain of PDS5C binds H3K4me1 and facilitates homology directed repair (HDR) (Niu et al. 2021). Other repair genes including PDS5 paralogs and MSH6 (responsible for mismatch repair − MMR) also contain functionally similar Tudor domains, with H3K4me1 targeting potential.

## Acknowledgments

The research was conducted at the University of California Davis, which is located on land which was the home of the Patwin people for thousands of years. We thank Satoyo Oya, Tetsuji Kakutani, and Soichi Inagaki for their feedback and insights on H3K4me1. The Monroe Lab is supported by FFAR grant ICRC20-0000000014, USDA-NIFA grant 108681-Z5327202, UC Davis STAIR grant, and CPRB grant HG-2022-35.

## Author Contributions

DQ, DL, and GM designed the research; DQ, DL, KZ, LO, AA and GM performed research; DQ, DL, KZ, AP, VY, PC, and GM contributed new analytic/computational/etc. tools; DQ, DL, KZ VY, AP, PC, and GM analyzed data; and DQ, DL, KZ, VY, PC, and GM wrote the paper.

## Methods

### Mutation dataset in *O. sativa*

Germline *de novo* mutations in 1,504 fast neutron mutagenesis lines were downloaded from Kitbase at kitbase.ucdavis.edu. These were independently called and validated as previously described (Li et al., 2017). We focused specifically on single base substitutions (SBS), which were validated with a >99% accuracy by Li et al. (2017). We annotated each SBS in coding regions as being a synonymous or non-synonymous mutation based on the effect on the amino acid sequence. We compared non-synonymous and synonymous ratios with values from genomes of 3,010 natural accessions (Wang et al., 2018) and neutral expectations based on mutation spectra, coding region nucleotide composition, and codon table with the *Null_ns_s* function from the *polymorphology* package in R (https://github.com/greymonroe/polymorphology/blob/main/R/Null_ns_s.R).

### Epigenomic data collection

Epigenome features were accessed from the RiceENCODE database (glab.hzau.edu.cn/RiceENCODE/), which has been previously described (Xie et al., 2021). In brief, peaks were called from ChIP-seq data with MACS2 (Zhang et al., 2008) narrow-peak calling settings. We analyzed peak distributions for H3K4me1, H3K9me1, H3K4me3, H3K36me3, H3K9me2, H3K27me3, H3K27ac, H3K4ac, H3K12ac, H3K9ac, and RNA polymerase II (PII) measured in Nipponbare *O. sativa* plant seedlings, which constituted the most complete set of histone modifications available. We repeated analyses but with H3K4me1 measurements derived from panicles and leaves (rather than seedlings) and found essentially the same results.

For downstream analyses comparing changes in mutation in *MSH6* knockout mutant lines as a function of H3K4me1, we accessed a collection of H3K4me1 ChIPseq datasets from *A. thaliana* maintained by the Plant Chromatin State Database (Liu et al., 2018). From these, we calculated the normalized mean across all datasets in 200 bp sliding (100 bp) windows across the entire genome.

### Estimation of the relationship between mutation rates and *O. sativa* epigenomic features

We divided the genome into 100 bp windows surrounding genes (+-3000 bp of genes). This allowed us to, in later steps, restrict our analyses to only genes known to have accumulated loss-of-function mutations, and thus be less likely to be affected by selection. We also divided the genome into 100bp windows and repeated analyses, to confirm that results were generally the same. We calculated the number of single base-pair substitutions and peaks for each epigenomic feature overlapping within each window. We then estimated the relationships between epigenomic features and mutation rates with a binomial generalized linear model where the response was a binary state defined as whether a substitution occurred in that window, predicted by all features, with predictors defined as whether that window overlapped with an epigenome peak. We also repeated the analyses with a linear regression model where the response was the number of mutations in a window and found essentially the same results, so we show the binomial regression results. To test whether findings were driven simply by GC>AT mutations in transposable elements, we removed all GC>AT and repeated analyses. To further control for any residual selection in the mutation accumulation experiment, we also restricted our analyses to genes harboring loss-of-function mutations in the population and repeated the analyses. Finally, we restricted analyses to homozygous SBS and repeated the analyses. Mutation frequencies were plotted around genes in 100 bp windows. Since gene bodies are different lengths, the position of the window was converted into a percent of gene length. H3K4me1 peaks around gene bodies were plotted similarly. We also visualized mutation frequencies relative to H3K4me1 peaks in the same manner.

### Analysis of ChIP-seq data of AtPDS5C

To study the distribution of PDS5C, we used ChIP-seq data as described by Niu et al. (2021). PDS5C enrichment was calculated as described by Niu et al. (2021) among regions as log2[(1 + n_ChIP)/N_ChIP] – log2[(1 + n_Input)/N_Input)], where n_ChIP and n_Input represent the total depth of mapped ChIP and Input fragments in a region, and N_ChIP and N_Input are the numbers total depths of mapped unique fragments. We calculated PDS5C enrichment in genic features (1000 bp upstream and downstream of genes, UTRs, introns, coding regions) and gene bodies (TSS to TTS) across the TAIR10 *A. thaliana* genome (arabidopsis.org).

### Relationship between AtPDS5C and functional constraint

We analyzed the enrichment of the PDS5C ChIP-seq peaks in *A. thaliana* in genetic features and estimated the relationships between those regions and mutation rates, H4K4me1, Pn/Ps, and tissue expression depth. Tissue expression data are from (Mergner et al., 2020). H3K4me1 in Arabidopsis is from the Plant Chromatin State Database (Liu et al., 2018). Synonymous (Ps) and non-synonymous polymorphism (Pn) data are from the 1001 Genomes project (1001 Genomes Consortium, 2016). Essential genes were based on findings from (Lloyd and Meinke, 2012). Germline mutation rates are from (Weng et al., 2019; Monroe et al., 2022).

### *Blastp* and protein structure prediction and visualization

We used *blastp* on Phytozome (Goodstein et al., 2012) to search the *O. sativa* proteome for PDS5C and MHS6 orthologs and to search the *A. thaliana* proteome for genes containing Tudor domains similar to that of PDS5C, which was validated experimentally to bind H3K4me1 (Niu et al., 2021). We submitted the resulting list of 29 genes with putative Tudor domains to gene ontology analysis with ShinyGO (bioinformatics.sdstate.edu/go/)(Ge et al., 2019).

Protein structure predictions were performed using AlphaFold (Jumper et al., 2021) in Google Colab (Mirdita et al., 2022) in no-template mode. All structures were visualized, processed, and analyzed using UCSF ChimeraX (Goddard et al., 2018).

### Peptide docking

H3 tail peptides comprising 5 amino acids with the different methylation states for K4 (none, mono, di or trimethylated) were docked to the experimental structure of *A. thaliana* PDS5C Tudor domain and to the models of *A. thaliana* and *O. sativa* MSH6 and *O. sativa* PDS5C Tudor domains using Rosetta FlexPepDock tool (Raveh et al., 2010) in refinement mode. We generated 10,000 docked models per case and analyzed the top 10% based on Rosetta’s total score. Analysis of the outputs was conducted using PyRosetta home-made scripts (Chaudhury et al., 2010) to measure the average distance of lysine 4 nitrogen in histone 3 to the center of the aromatic rings in the residues forming the aromatic cage in Tudor domains.

### Somatic mutations in *MSH6* Knockout lines

To study the effect of MSH6 in H3K4me1 targeted repair we performed 250x coverage sequencing analyses of 17 *A. thaliana*; 9 *WT/WT*, 7 *msh6/msh6* (6 SALK_037557 and 1 SALK_089638) and 2 *msh6/WT* (2 heterozygous SALK_089638). In brief, 3 week old plants were harvested, stems and roots were removed and rosettes were stored at -80 ºC. Whole rosettes were grinded using a mortar and liquid nitrogen and Qiagen DNeasy Plant Mini Kit (Cat No: 69106) following the manufacturer’s instructions. Illumina library preparation and Whole-genome DNA sequencing with NovaSeq (150-bp read length at high genomic coverage ∼250x) was performed by the UC Davis Genome Center. Reads were trimmed with trimmomatic (Bolger et al., 2014) and duplicates were marked with the samtools markdup function (Li et al., 2009). Reads were mapped to the *A. thaliana* TAIR10 reference genome with bwa mem(Li and Durbin, 2009). Strelka2 variant caller in TUMOR-NORMAL mode (each sample was compared to all other 16 samples as NORMALs) was used to call SBS in the sequenced samples (Kim et al., 2018). Mutations found were filtered by keeping variants called in only one sample, given a PASS quality score in all 16 “NORMAL” comparisons. In addition to the built-in filters imposed by Strelka2 for variants to receive a PASS, which “include (1) the genotype probability computed by the core variant probability model, (2) root-mean-square mapping quality, (3) strand bias, (4) the fraction of reads consistent with locus haplotype model, and (5) the complexity of the reference context as measured by metrics such as homopolymer length and compressibility” (Kim et al., 2018), we further filtered to only include variants in which the distance to the nearest mutation was >1000 bp, total read depth >150 and alt read support > 5. To distinguish somatic mutations from de novo heterozygous variants, read-depths between alternative and reference alleles were compared with an *X*^2^ test against the expected 50/50 ratio for heterozygous variants. Mutations were considered somatic if the X^2^ test statistic *p-value* was less than 0.001 (to account for multiple testing) and the alt read depth was < 50% Homozygous *de novo* germline mutations were identified when X^2^ test statistic p-value was less than 0.001 and the alt read depth was >50%. Complete code for variant calling, post-processing, and analyses are maintained on https://github.com/greymonroe/Quiroz_H3K4me1_mediated_repair.

### Code and data

Figures, code, and data are located on: https://github.com/greymonroe/Quiroz_H3K4me1_mediated_repair.

---

**Figure S1.**
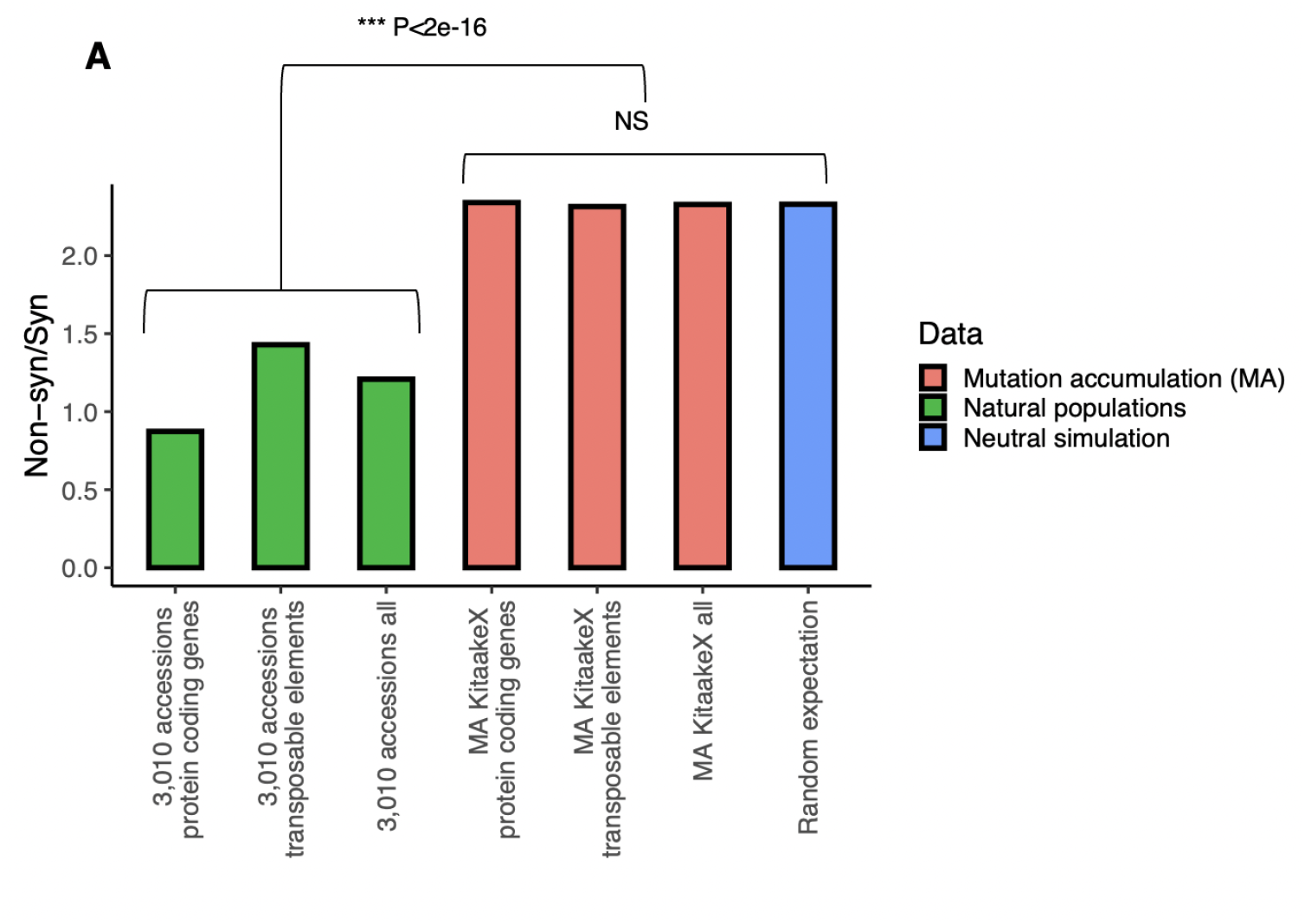
Non-synonymous to synonymous ratios. The non-synonymous to synonymous ratio was significantly lower (p<2.×10^−16^) in natural populations compared to mutation accumulation lines and neutral expectation. The non-synonymous to synonymous ratio was similar in mutation accumulation lines to neutral expectation.

**Figure S2.**
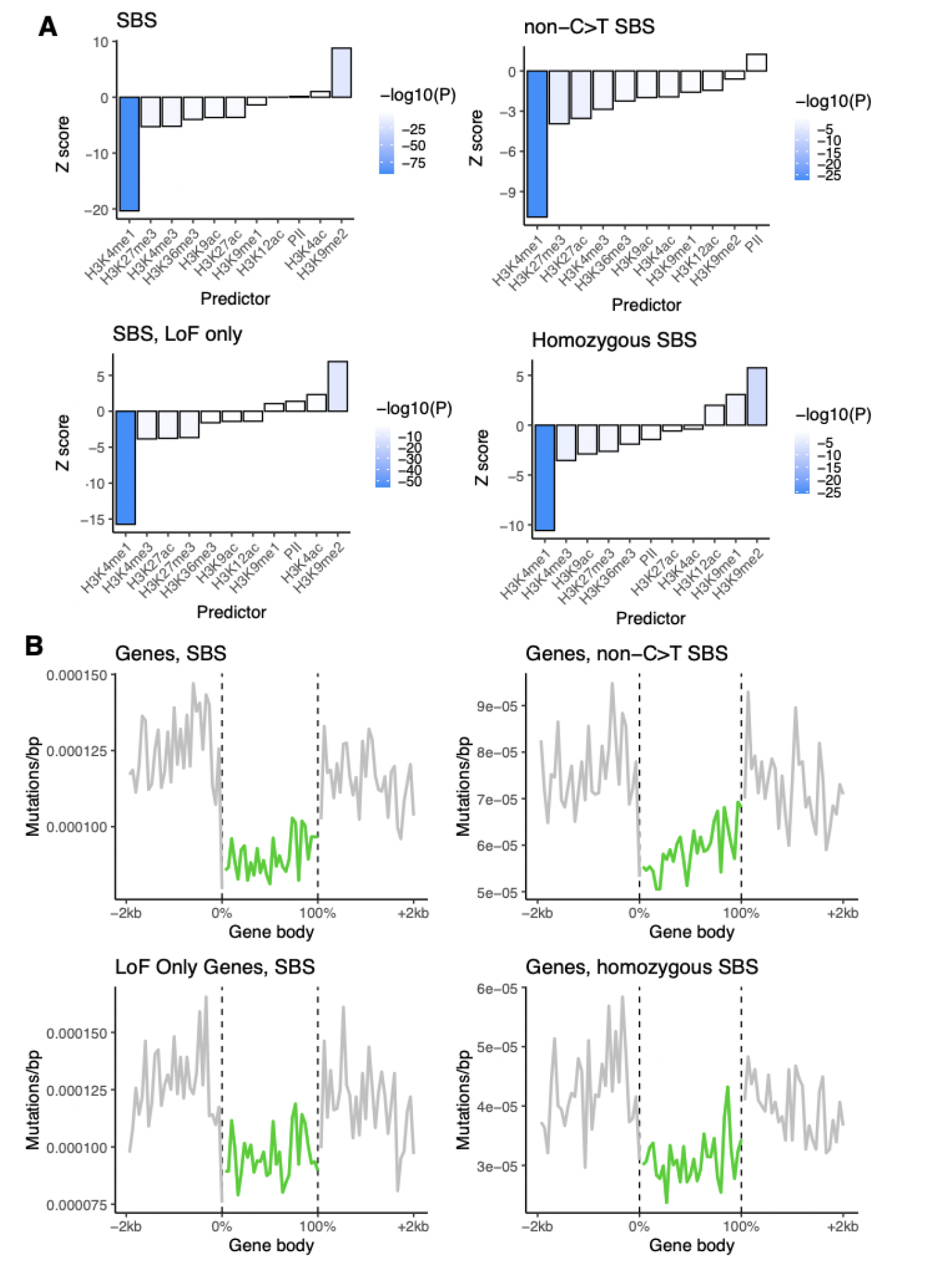
Relationships between epigenomic features, mutation and gene bodies. **A**, Results from logistic regression modeling mutation probability in 100bp windows as a function of overlap with epigenomic marks around genes in rice. Four panels reflect various subsets of data to test for effects of cytosine deamination in transposable elements as alternative explanation, selection, or mapping. **B**, Mutation rates in relation to gene bodies in rice.

**Figure S3.**
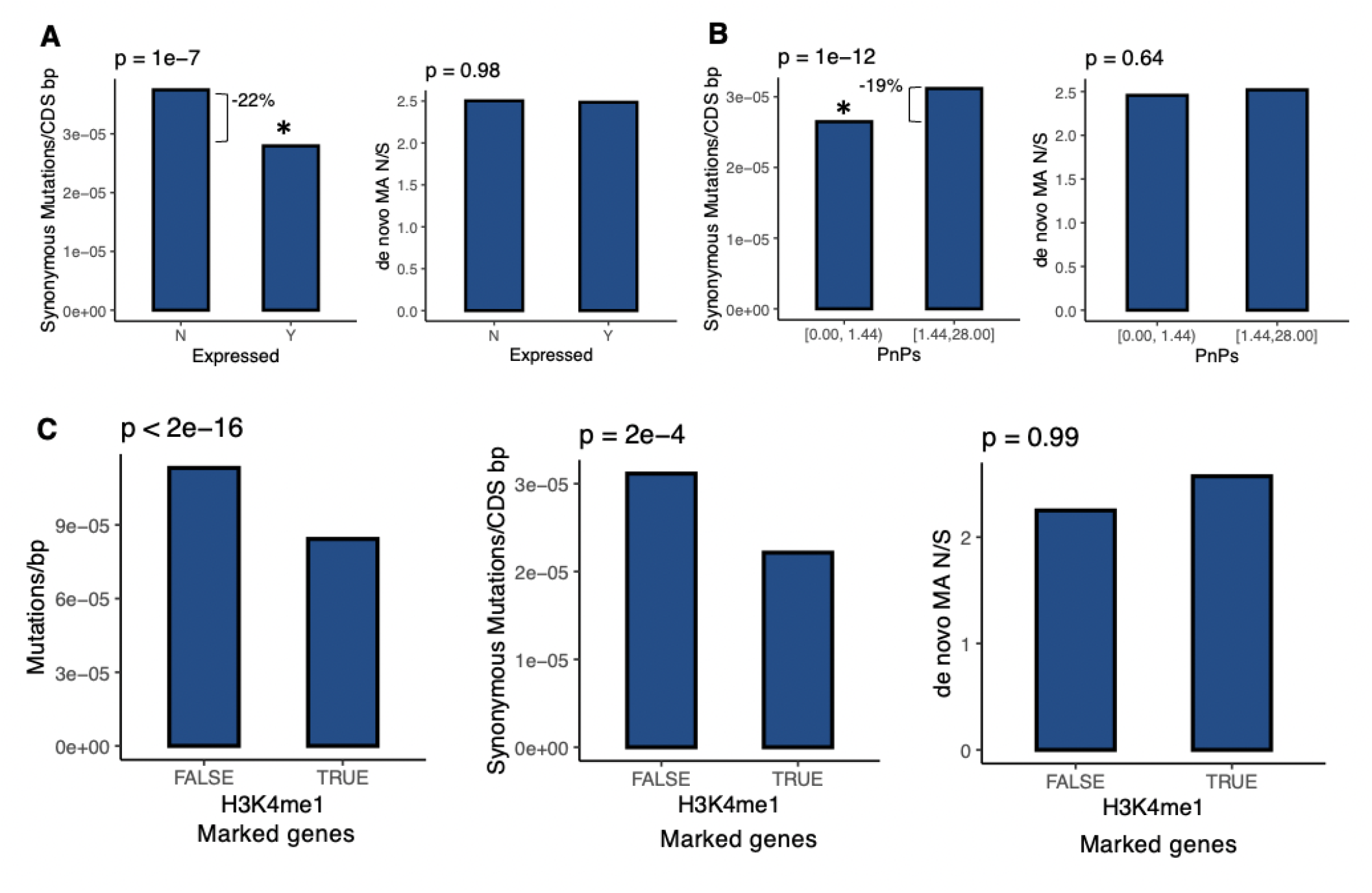
Expressed and conserved genes and H3K4me1 marked genes have lower mutation rates in rice. **A**, synonymous mutation rates, non-synonymous to synonymous *de novo* mutation ratio in mutation accumulation lines (MA N/S) in expressed vs non-expressed genes and **B**, in genes subject variable degrees of purifying selection (low vs high Pn/Ps) in 3,010 natural accessions of rice. **C**, mutation rates, non-synonymous to synonymous *de novo* mutations in mutation accumulation lines (MA N/S) in H3K4me1 non-marked genes (FALSE) and marked genes (TRUE).

**Figure S4.**
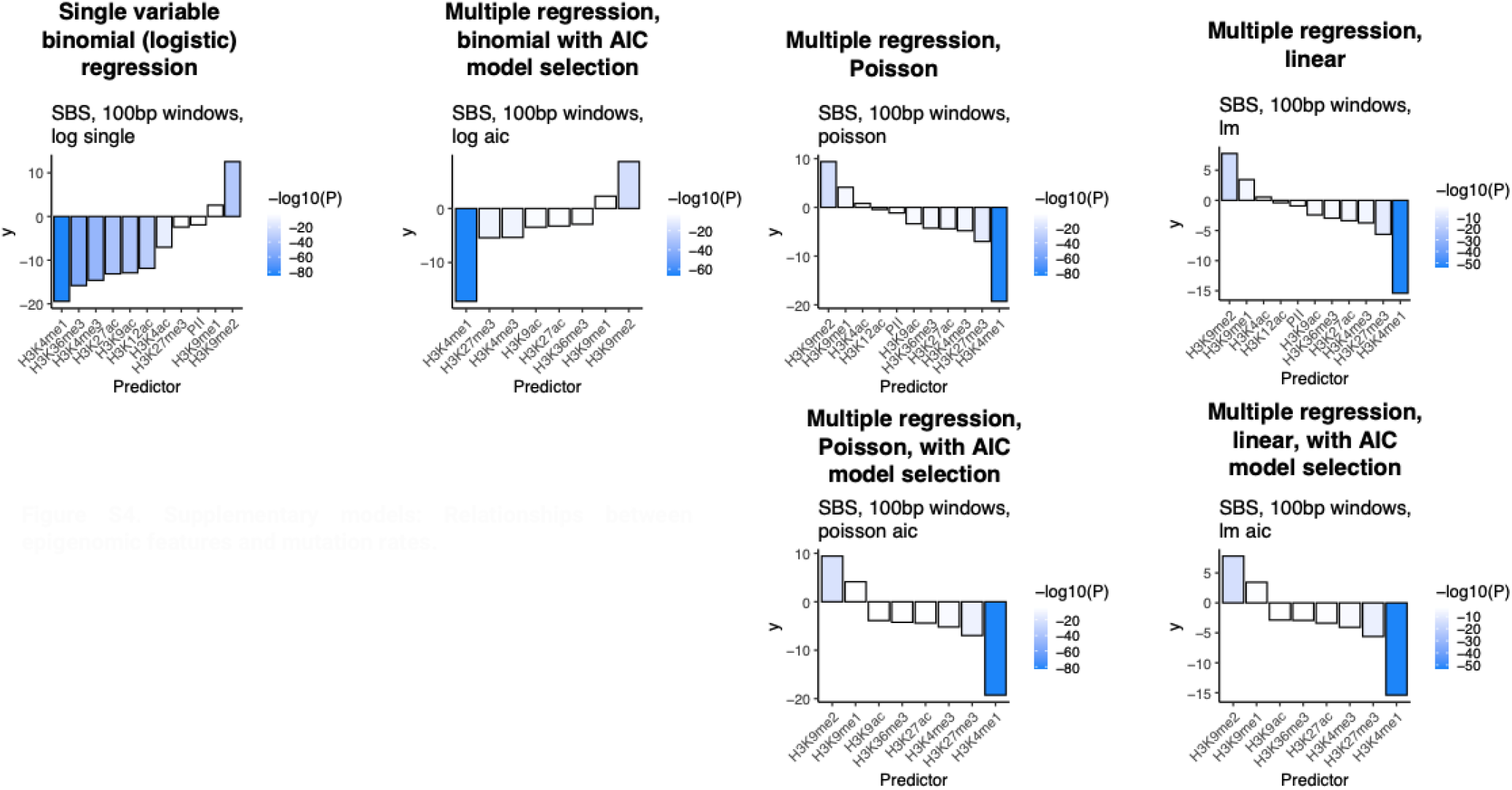
Supplementary models: Relationships between epigenomic features and mutation rates.

**Figure S5.**
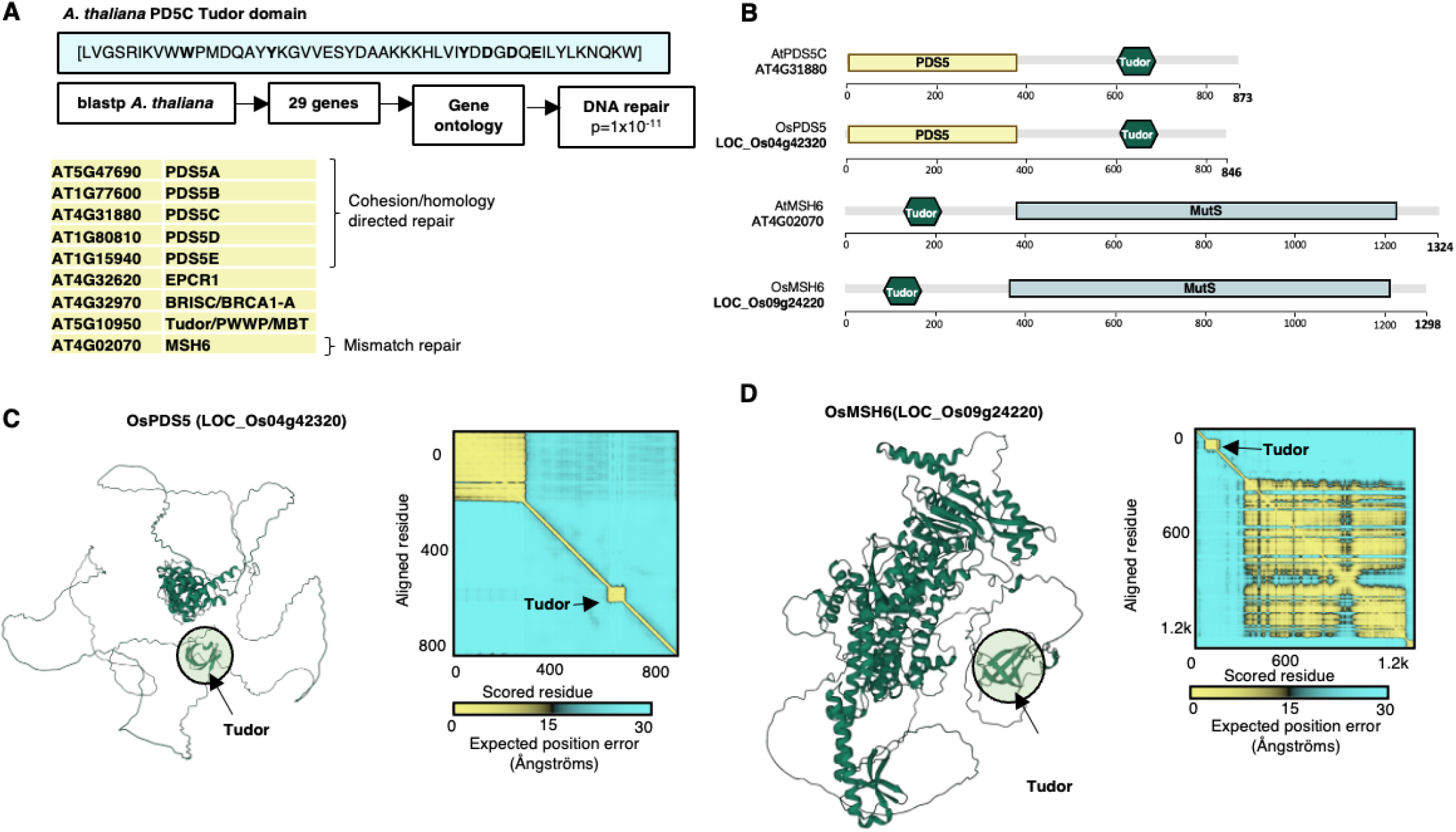
Multiple repair pathways with H3K4me1 targeting potential via Tudor domains. **A**, Results from blastp of PCDS5C Tudor domain, which has confirmed H3K4me1 binding specificity (Niu et al. 2021) identified 29 genes. Nine of these are annotated as having DNA repair function (significant enrichment by gene ontology analysis)., **B**, Domain prediction of *A. thaliana* and *O. sativa* putative orthologs of PDS5C and MSH6 genes. **C-D**, Structure of PDS5C and MSH6 protein from Alphafold2 indicates Tudor domain is tethered to active domains structure in putative *O. sativa* orthologs.

**Figure S6.**
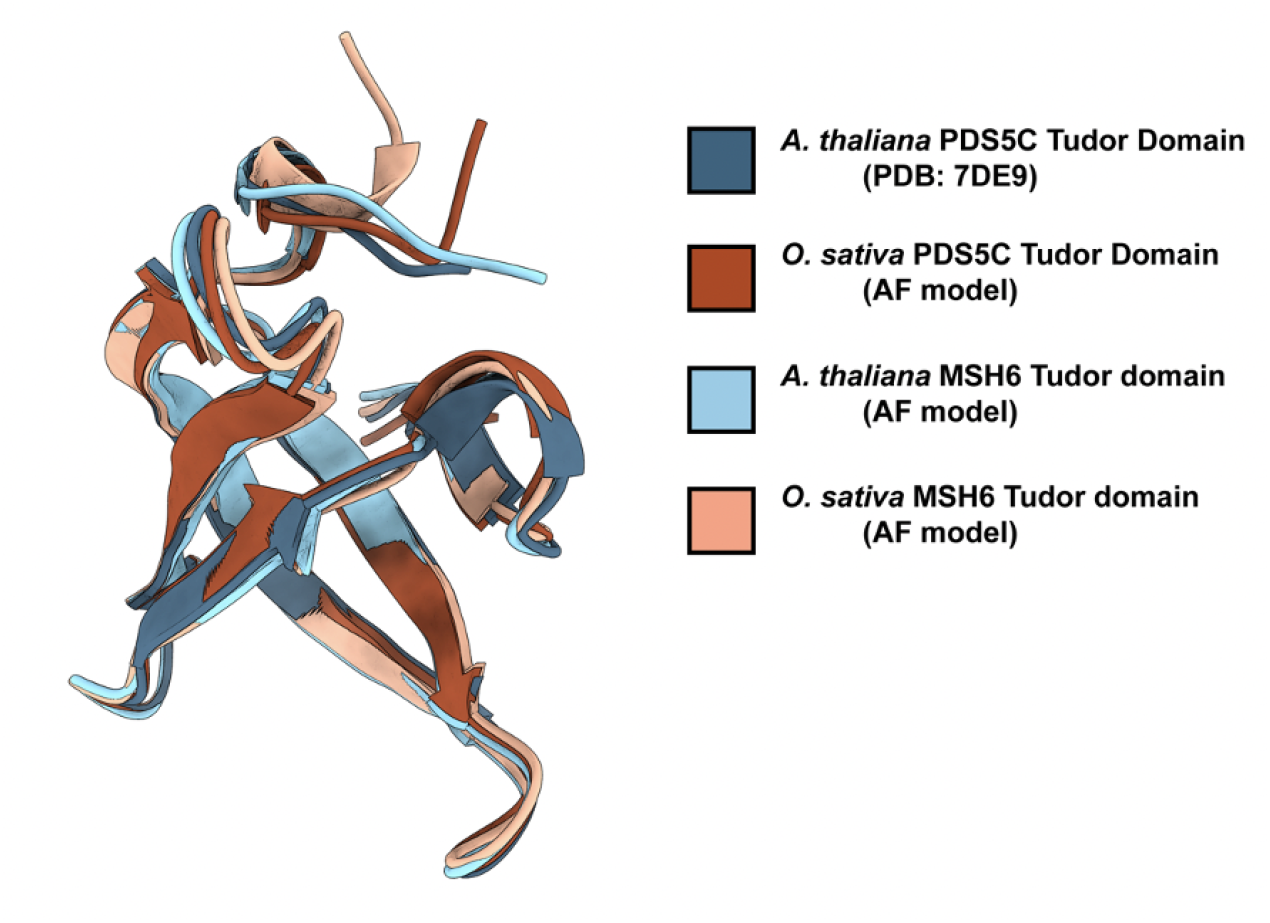
AlphaFold predictions of Tudor domains. AlphaFold models of *A. thaliana* and *O. sativa MSH6* and *O. sativa* PDS5C Tudor are shown superimposed onto the experimental structure of A. thaliana PDS5C Tudor domain (PDB:7DE9) (Niu et al. 2021).

**Figure S7.**
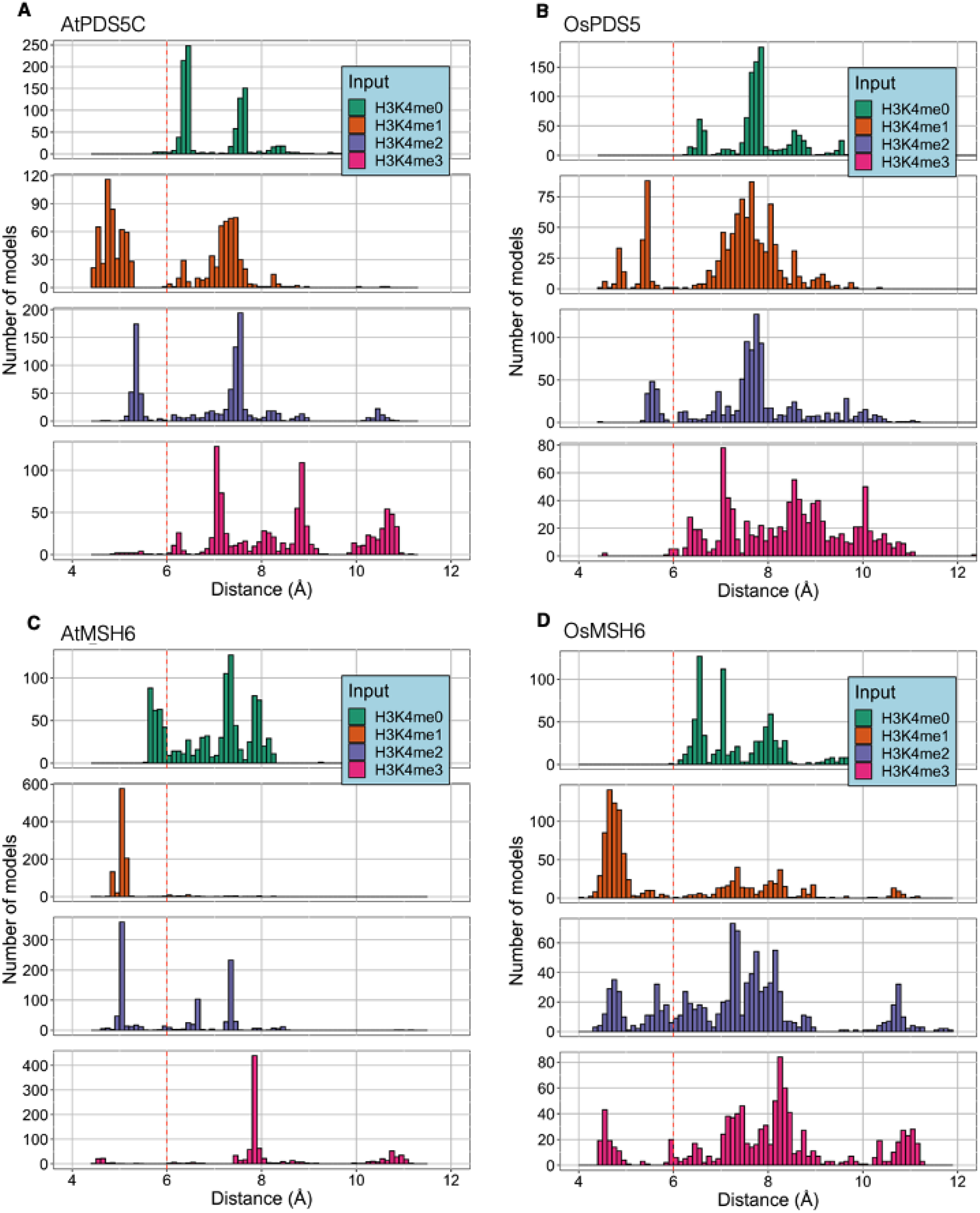
The average distance (In Å) distribution to the aromatic cage In FlexPepDock models of the different H3K4 marks tested, for; **A**, *A. thaliana* PDS5C Tudor experimental structure (PDB:7de9) **B**, *O. sativa* PDS5C Tudor Alphafold model **C**, *A. thaliana* MSH6 Tudor Alphafold model D, *O. sativa* MSH6 Tudor Alphafold model. **A-D**, The red line marks the threshold of 6Å (defining the H3K4 inside the aromatic cage).

**Figure S8.**
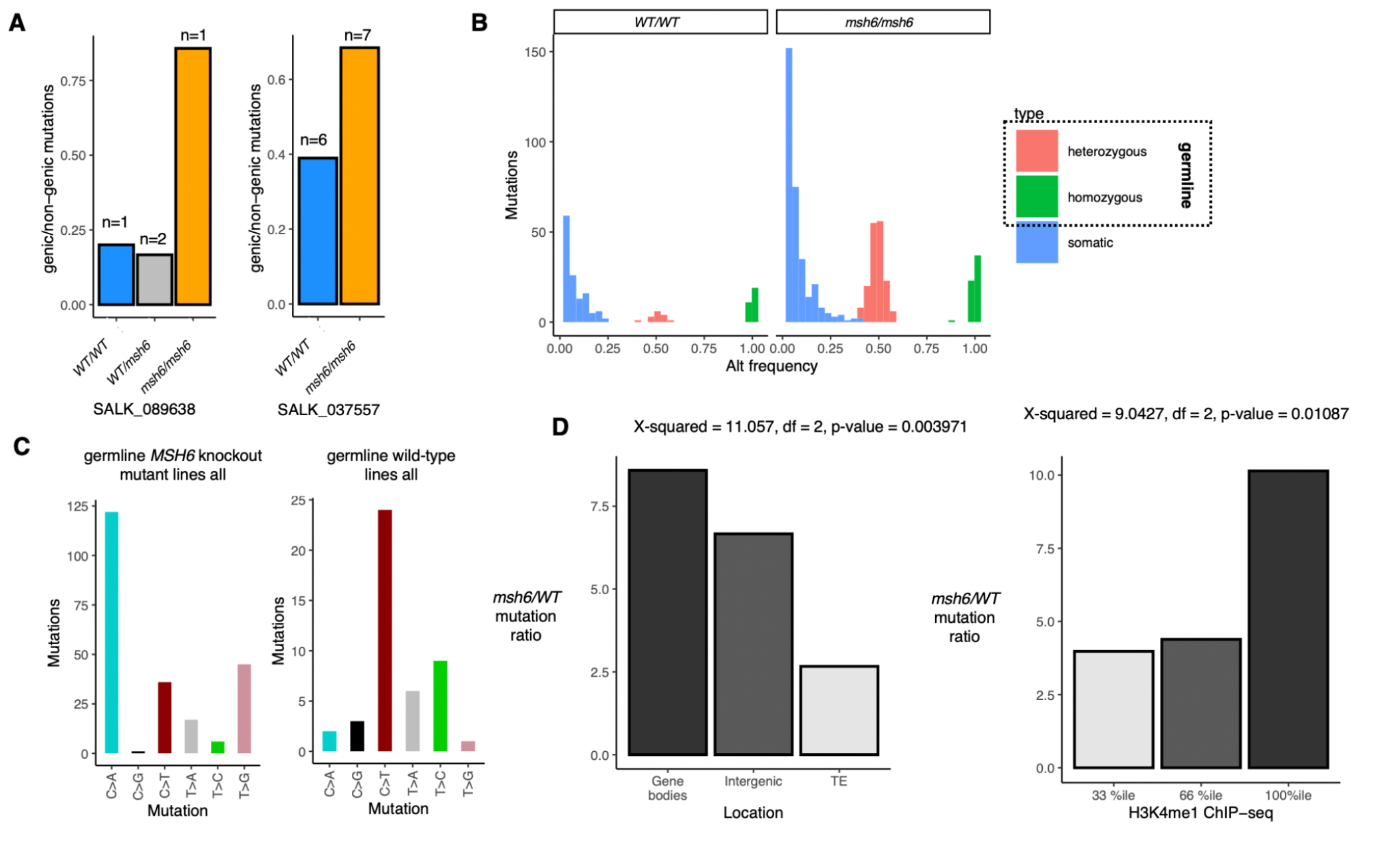
**A**, genic/non-genic mutations in *MSH6* knockout mutant lines SALK_089638 and SALK_037557. **B**, frequency of alternative alleles distribution detected by Strelka2 variant caller. **C**, Germline mutations signature in *MSH6* knockout mutant lines and wild-type lines. **D**, *msh6/WT* mutation ratio in different genomic features and in different H3K4me1 ChlP-seq %ile for *de novo* germline mutations.

**Supplemental Table 1.**
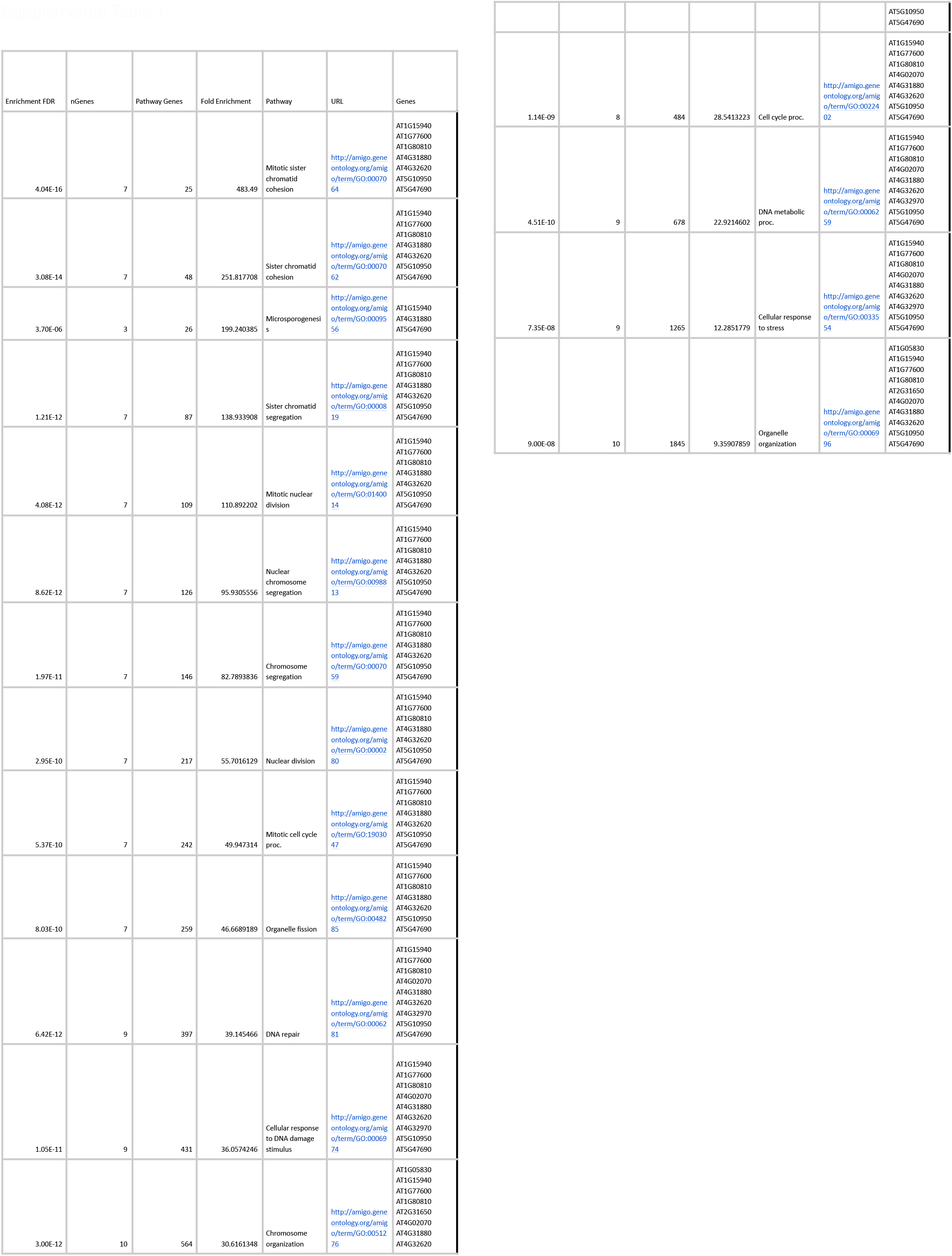

## References

1001 Genomes Consortium (2016). 1,135 Genomes Reveal the Global Pattern of Polymorphism in Arabidopsis thaliana. Cell 166: 481–491.

Adé, J., Belzile, F., Philippe, H., and Doutriaux, M.P. (1999). Four mismatch repair paralogues coexist in Arabidopsis thaliana: AtMSH2, AtMSH3, AtMSH6-1 and AtMSH6-2. Mol. Gen. Genet. 262: 239–249.

Akdemir, K.C. et al. (2020). Somatic mutation distributions in cancer genomes vary with three-dimensional chromatin structure. Nat. Genet. 52: 1178–1188.

Aymard, F., Bugler, B., Schmidt, C.K., Guillou, E., Caron, P., Briois, S., Iacovoni, J.S., Daburon, V., Miller, K.M., Jackson, S.P., and Legube, G. (2014). Transcriptionally active chromatin recruits homologous recombination at DNA double-strand breaks. Nat. Struct. Mol. Biol. 21: 366–374.

Belfield, E.J., Brown, C., Ding, Z.J., Chapman, L., Luo, M., Hinde, E., van Es, S.W., Johnson, S., Ning, Y., Zheng, S.J., Mithani, A., and Harberd, N.P. (2021). Thermal stress accelerates Arabidopsis thaliana mutation rate. Genome Res. 31: 40–50.

Belfield, E.J., Ding, Z.J., Jamieson, F.J.C., Visscher, A.M., Zheng, S.J., Mithani, A., and Harberd, N.P. (2018). DNA mismatch repair preferentially protects genes from mutation. Genome Res. 28: 66–74.

Bolaños-Villegas, P., Yang, X., Wang, H.-J., Juan, C.-T., Chuang, M.-H., Makaroff, C.A., and Jauh, G.-Y. (2013). Arabidopsis CHROMOSOME TRANSMISSION FIDELITY 7 (AtCTF7/ECO1) is required for DNA repair, mitosis and meiosis. Plant J. 75: 927–940.

Bolger, A.M., Lohse, M., and Usadel, B. (2014). Trimmomatic: a flexible trimmer for Illumina sequence data. Bioinformatics 30: 2114–2120.

Cagan, A. et al. (2022). Somatic mutation rates scale with lifespan across mammals. Nature 604: 517–524.

Chaudhury, S., Lyskov, S., and Gray, J.J. (2010). PyRosetta: a script-based interface for implementing molecular modeling algorithms using Rosetta. Bioinformatics 26: 689–691.

Davarinejad, H. et al. (2022). The histone H3.1 variant regulates TONSOKU-mediated DNA repair during replication. Science 375: 1281–1286.

Fang, H., Zhu, X., Yang, H., Oh, J., Barbour, J.A., and Wong, J.W.H. (2021). Deficiency of replication-independent DNA mismatch repair drives a 5-methylcytosine deamination mutational signature in cancer. Sci Adv 7: eabg4398.

Foster, P.L., Lee, H., Popodi, E., Townes, J.P., and Tang, H. (2015). Determinants of spontaneous mutation in the bacterium Escherichia coli as revealed by whole-genome sequencing. Proc. Natl. Acad. Sci. U. S. A. 112: E5990–9.

Ge, S.X., Jung, D., and Yao, R. (2019). ShinyGO: a graphical gene-set enrichment tool for animals and plants. Bioinformatics 36: 2628–2629.

Goddard, T.D., Huang, C.C., Meng, E.C., Pettersen, E.F., Couch, G.S., Morris, J.H., and Ferrin, T.E. (2018). UCSF ChimeraX: Meeting modern challenges in visualization and analysis. Protein Sci. 27: 14–25.

Gonzalez, V. and Spampinato, C.P. (2020). The mismatch repair protein MSH6 regulates somatic recombination in Arabidopsis thaliana. DNA Repair 87: 102789.

Goodstein, D.M., Shu, S., Howson, R., Neupane, R., Hayes, R.D., Fazo, J., Mitros, T., Dirks, W., Hellsten, U., Putnam, N., and Rokhsar, D.S. (2012). Phytozome: a comparative platform for green plant genomics. Nucleic Acids Res. 40: D1178–86.

Gu, Y., Parker, A., Wilson, T.M., Bai, H., Chang, D.-Y., and Lu, A.-L. (2002). Human MutY Homolog, a DNA Glycosylase Involved in Base Excision Repair, Physically and Functionally Interacts with Mismatch Repair Proteins Human MutS Homolog 2/Human MutS Homolog 6*. J. Biol. Chem. 277: 11135–11142.

Habig, M., Lorrain, C., Feurtey, A., Komluski, J., and Stukenbrock, E.H. (2021). Epigenetic modifications affect the rate of spontaneous mutations in a pathogenic fungus. Nat. Commun. 12: 5869.

Hahm, J.Y., Park, J., Jang, E.-S., and Chi, S.W. (2022). 8-Oxoguanine: from oxidative damage to epigenetic and epitranscriptional modification. Exp. Mol. Med. 54: 1626–1642.

Hill, V.K., Kim, J.-S., and Waldman, T. (2016). Cohesin mutations in human cancer. Biochim. Biophys. Acta 1866: 1–11.

Huang, Y., Gu, L., and Li, G.-M. (2018). H3K36me3-mediated mismatch repair preferentially protects actively transcribed genes from mutation. J. Biol. Chem. 293: 7811–7823.

Jiang, P., Ollodart, A.R., Sudhesh, V., Herr, A.J., Dunham, M.J., and Harris, K. (2021). A modified fluctuation assay reveals a natural mutator phenotype that drives mutation spectrum variation within Saccharomyces cerevisiae. Elife 10.

Jumper, J. et al. (2021). Highly accurate protein structure prediction with AlphaFold. Nature 596: 583–589.

Katju, V., Konrad, A., Deiss, T.C., and Bergthorsson, U. (2022). Mutation rate and spectrum in obligately outcrossing Caenorhabditis elegans mutation accumulation lines subjected to RNAi-induced knockdown of the mismatch repair gene msh-2. G3 12: jkab364.

Kawahara, Y. et al. (2013). Improvement of the Oryza sativa Nipponbare reference genome using next generation sequence and optical map data. Rice 6: 4.

Kim, J., Daniel, J., Espejo, A., Lake, A., Krishna, M., Xia, L., Zhang, Y., and Bedford, M.T. (2006). Tudor, MBT and chromo domains gauge the degree of lysine methylation. EMBO Rep. 7: 397–403.

Kim, S., Scheffler, K., Halpern, A.L., Bekritsky, M.A., Noh, E., Källberg, M., Chen, X., Kim, Y., Beyter, D., Krusche, P., and Saunders, C.T. (2018). Strelka2: fast and accurate calling of germline and somatic variants. Nat. Methods 15: 591–594.

Kino, K., Hirao-Suzuki, M., Morikawa, M., Sakaga, A., and Miyazawa, H. (2017). Generation, repair and replication of guanine oxidation products. Genes Environ 39: 21.

Kolodner, R. (1996). Biochemistry and genetics of eukaryotic mismatch repair. Genes Dev. 10: 1433–1442.

Krasovec, M., Eyre-Walker, A., Sanchez-Ferandin, S., and Piganeau, G. (2017). Spontaneous Mutation Rate in the Smallest Photosynthetic Eukaryotes. Mol. Biol. Evol. 34: 1770–1779.

Li, F., Mao, G., Tong, D., Huang, J., Gu, L., Yang, W., and Li, G.-M. (2013). The histone mark H3K36me3 regulates human DNA mismatch repair through its interaction with MutSα. Cell 153: 590–600.

Li, G. et al. (2017). The Sequences of 1504 Mutants in the Model Rice Variety Kitaake Facilitate Rapid Functional Genomic Studies. Plant Cell 29: 1218–1231.

Li, H. and Durbin, R. (2009). Fast and accurate short read alignment with Burrows–Wheeler transform. Bioinformatics 25: 1754–1760.

Li, H., Handsaker, B., Wysoker, A., Fennell, T., Ruan, J., Homer, N., Marth, G., Abecasis, G., and Durbin, R. (2009). The Sequence Alignment/Map format and SAMtools. Bioinformatics 25: 2078–2079.

Li, L., Jean, M., and Belzile, F. (2006). The impact of sequence divergence and DNA mismatch repair on homeologous recombination in Arabidopsis. Plant J. 45: 908–916.

Li, R. et al. (2021). A body map of somatic mutagenesis in morphologically normal human tissues. Nature 597: 398–403.

Liu, H. and Zhang, J. (2022). Is the Mutation Rate Lower in Genomic Regions of Stronger Selective Constraints? Mol. Biol. Evol. 39.

Liu, Q., Liu, P., Ji, T., Zheng, L., Shen, C., Ran, S., Liu, J., Zhao, Y., Niu, Y., Wang, T., and Dong, J. (2022). The histone methyltransferase SUVR2 promotes DSB repair via chromatin remodeling and liquid-liquid phase separation. Mol. Plant.

Liu, Y., Tian, T., Zhang, K., You, Q., Yan, H., Zhao, N., Yi, X., Xu, W., and Su, Z. (2018). PCSD: a plant chromatin state database. Nucleic Acids Res. 46: D1157–D1167.

Lloyd, J. and Meinke, D. (2012). A comprehensive dataset of genes with a loss-of-function mutant phenotype in Arabidopsis. Plant Physiol. 158: 1115–1129.

Lloyd, J.P., Seddon, A.E., Moghe, G.D., Simenc, M.C., and Shiu, S.-H. (2015). Characteristics of Plant Essential Genes Allow for within- and between-Species Prediction of Lethal Mutant Phenotypes. Plant Cell 27: 2133–2147.

López-Cortegano, E., Craig, R.J., Chebib, J., Samuels, T., Morgan, A.D., Kraemer, S.A., Böndel, K.B., Ness, R.W., Colegrave, N., and Keightley, P.D. (2021). De Novo Mutation Rate Variation and Its Determinants in Chlamydomonas. Mol. Biol. Evol. 38: 3709–3723.

Lujan, S.A., Clausen, A.R., Clark, A.B., MacAlpine, H.K., MacAlpine, D.M., Malc, E.P., Mieczkowski, P.A., Burkholder, A.B., Fargo, D.C., Gordenin, D.A., and Kunkel, T.A. (2014). Heterogeneous polymerase fidelity and mismatch repair bias genome variation and composition. Genome Res. 24: 1751.

Lu, R. and Wang, G.G. (2013). Tudor: a versatile family of histone methylation “readers.” Trends Biochem. Sci. 38: 546–555.

Lu, Z., Cui, J., Wang, L., Teng, N., Zhang, S., Lam, H.-M., Zhu, Y., Xiao, S., Ke, W., Lin, J., Xu, C., and Jin, B. (2021). Genome-wide DNA mutations in Arabidopsis plants after multigenerational exposure to high temperatures. Genome Biol. 22: 160.

Lynch, M. (2010). Evolution of the mutation rate. Trends Genet. 26: 345–352.

Lynch, M., Ackerman, M.S., Gout, J.-F., Long, H., Sung, W., Thomas, W.K., and Foster, P.L. (2016). Genetic drift, selection and the evolution of the mutation rate. Nat. Rev. Genet. 17: 704–714.

Makova, K.D. and Hardison, R.C. (2015). The effects of chromatin organization on variation in mutation rates in the genome. Nat. Rev. Genet. 16: 213–223.

Martincorena, I. and Luscombe, N.M. (2013). Non-random mutation: the evolution of targeted hypermutation and hypomutation. Bioessays 35: 123–130.

Maurer-Stroh, S., Dickens, N.J., Hughes-Davies, L., Kouzarides, T., Eisenhaber, F., and Ponting, C.P. (2003). The Tudor domain “Royal Family”: Tudor, plant Agenet, Chromo, PWWP and MBT domains. Trends Biochem. Sci. 28: 69–74.

Mazurek, A., Berardini, M., and Fishel, R. (2002). Activation of Human MutS Homologs by 8-Oxo-guanine DNA Damage*. J. Biol. Chem. 277: 8260–8266.

Mergner, J. et al. (2020). Mass-spectrometry-based draft of the Arabidopsis proteome. Nature 579: 409–414.

Mirdita, M., Schütze, K., Moriwaki, Y., Heo, L., Ovchinnikov, S., and Steinegger, M. (2022). ColabFold: making protein folding accessible to all. Nat. Methods 19: 679–682.

Monroe, J.G. et al. (2022). Mutation bias reflects natural selection in Arabidopsis thaliana. Nature.

Moore, L. et al. (2021). The mutational landscape of human somatic and germline cells. Nature.

Morales, C., Ruiz-Torres, M., Rodríguez-Acebes, S., Lafarga, V., Rodríguez-Corsino, M., Megías, D., Cisneros, D.A., Peters, J.-M., Méndez, J., and Losada, A. (2020). PDS5 proteins are required for proper cohesin dynamics and participate in replication fork protection. J. Biol. Chem. 295: 146–157.

Ni, T.T., Marsischky, G.T., and Kolodner, R.D. (1999). MSH2 and MSH6 are required for removal of adenine misincorporated opposite 8-oxo-guanine in S. cerevisiae.Mol. Cell 4: 439–444.

Niu, Q. et al. (2021). A histone H3K4me1-specific binding protein is required for siRNA accumulation and DNA methylation at a subset of loci targeted by RNA-directed DNA methylation. Nat. Commun. 12: 3367.

Oya, S., Takahashi, M., Takashima, K., Kakutani, T., and Inagaki, S. (2021). Transcription-coupled and epigenome-encoded mechanisms direct H3K4 methylation. bioRxiv: 2021.06.03.446702.

Pavlov, Y.I., Mian, I.M., and Kunkel, T.A. (2003). Evidence for preferential mismatch repair of lagging strand DNA replication errors in yeast. Curr. Biol. 13: 744–748.

de la Peña, M.V., Summanen, P.A.M., Liukkonen, M., and Kronholm, I. (2022). Chromatin structure influences rate and spectrum of spontaneous mutations in Neurospora crassa. bioRxiv: 2022.03.13.484164.

Phipps, J. and Dubrana, K. (2022). DNA Repair in Space and Time: Safeguarding the Genome with the Cohesin Complex. Genes 13.

Pradillo, M., Knoll, A., Oliver, C., Varas, J., Corredor, E., Puchta, H., and Santos, J.L. (2015). Involvement of the Cohesin Cofactor PDS5 (SPO76) During Meiosis and DNA Repair in Arabidopsis thaliana. Front. Plant Sci. 6: 1034.

Raveh, B., London, N., and Schueler-Furman, O. (2010). Sub-angstrom modeling of complexes between flexible peptides and globular proteins. Proteins 78: 2029–2040.

Ren, Q., Yang, H., Rosinski, M., Conrad, M.N., Dresser, M.E., Guacci, V., and Zhang, Z. (2005). Mutation of the cohesin related gene PDS5 causes cell death with predominant apoptotic features in Saccharomyces cerevisiae during early meiosis. Mutat. Res. 570: 163–173.

Sanders, M.A. et al. (2021). Life without mismatch repair. bioRxiv: 2021.04.14.437578.

Sasani, T.A., Ashbrook, D.G., Beichman, A.C., Lu, L., Palmer, A.A., Williams, R.W., Pritchard, J.K., and Harris, K. (2022). A natural mutator allele shapes mutation spectrum variation in mice. Nature 605: 497–502.

Schep, R. et al. (2021). Impact of chromatin context on Cas9-induced DNA double-strand break repair pathway balance. Mol. Cell 81: 2216–2230.e10.

Schubert, V., Weissleder, A., Ali, H., Fuchs, J., Lermontova, I., Meister, A., and Schubert, I. (2009). Cohesin gene defects may impair sister chromatid alignment and genome stability in Arabidopsis thaliana. Chromosoma 118: 591–605.

Schuster-Böckler, B. and Lehner, B. (2012). Chromatin organization is a major influence on regional mutation rates in human cancer cells. Nature 488: 504–507.

Sun, Z., Zhang, Y., Jia, J., Fang, Y., Tang, Y., Wu, H., and Fang, D. (2020). H3K36me3, message from chromatin to DNA damage repair. Cell Biosci. 10: 9.

Supek, F. and Lehner, B. (2017). Clustered Mutation Signatures Reveal that Error-Prone DNA Repair Targets Mutations to Active Genes. Cell 170: 534–547.e23.

Supek, F. and Lehner, B. (2015). Differential DNA mismatch repair underlies mutation rate variation across the human genome. Nature 521: 81–84.

Supek, F. and Lehner, B. (2019). Scales and mechanisms of somatic mutation rate variation across the human genome. DNA Repair 81: 102647.

Wang, W. et al. (2018). Genomic variation in 3,010 diverse accessions of Asian cultivated rice. Nature 557: 43–49.

Weiss, T., Crisp, P.A., Rai, K.M., Song, M., Springer, N.M., and Zhang, F. (2022). Drastic differential CRISPR-Cas9 induced mutagenesis influenced by DNA methylation and chromatin features. bioRxiv: 2022.02.28.482333.

Weng, M.-L., Becker, C., Hildebrandt, J., Neumann, M., Rutter, M.T., Shaw, R.G., Weigel, D., and Fenster, C.B. (2019). Fine-Grained Analysis of Spontaneous Mutation Spectrum and Frequency in Arabidopsis thaliana. Genetics 211: 703–714.

Wyant, S.R., Rodriguez, M.F., Carter, C.K., Parrott, W.A., Jackson, S.A., Stupar, R.M., and Morrell, P.L. (2022). Fast neutron mutagenesis in soybean enriches for small indels and creates frameshift mutations. G3 12.

Xie, L., Liu, M., Zhao, L., Cao, K., Wang, P., Xu, W., Sung, W.-K., Li, X., and Li, G. (2021). RiceENCODE: A comprehensive epigenomic database as a rice Encyclopedia of DNA Elements. Mol. Plant 14: 1604–1606.

Yang, X. et al. (2021). Developmental and temporal characteristics of clonal sperm mosaicism. Cell 184: 4772–4783.e15.

Yan, W., Deng, X.W., Yang, C., and Tang, X. (2021). The Genome-Wide EMS Mutagenesis Bias Correlates With Sequence Context and Chromatin Structure in Rice. Front. Plant Sci. 12: 579675.

Zhang, Y., Liu, T., Meyer, C.A., Eeckhoute, J., Johnson, D.S., Bernstein, B.E., Nusbaum, C., Myers, R.M., Brown, M., Li, W., and Liu, X.S. (2008). Model-based analysis of ChIP-Seq (MACS). Genome Biol. 9: R137.

Zhu, X., Xie, S., Tang, K., Kalia, R.K., Liu, N., Ma, J., Bressan, R.A., and Zhu, J.-K. (2021). Non-CG DNA methylation-deficiency mutations enhance mutagenesis rates during salt adaptation in cultured Arabidopsis cells. Stress Biology 1: 12.

